# Horizontal Transfer and Evolution of the Biosynthetic Gene Cluster for Benzoxazinoid

**DOI:** 10.1101/2021.12.07.471670

**Authors:** Dongya Wu, Bowen Jiang, Chu-Yu Ye, Michael P. Timko, Longjiang Fan

**Affiliations:** Institute of Crop Science & Institute of Bioinformatics, Zhejiang University, Hangzhou 310058, China; Department of Biology, University of Virginia, Charlottesville, VA 22904; Zhejiang University City College, Hangzhou 310015, China

**Keywords:** biosynthetic gene cluster, horizontal transfer, benzoxazinoid, grass, purifying selection

## Abstract

Benzoxazinoids are a class of protective and allelopathic plant secondary metabolites, first identified in maize (*Zea mays*) and subsequently shown to be encoded by a biosynthetic gene cluster (BGC), the Bx cluster. Data mining of mining 40 high-quality grass genomes identified complete Bx clusters (containing genes *Bx1* to *Bx5* and *Bx8*) in three genera (*Zea, Echinochloa* and *Dichanthelium*) in the Panicoideae and partial clusters in the Triticeae. The Bx cluster originated from gene duplication of native analogues of Bx genes and chromosomal translocation. An ancient Bx cluster including additional Bx genes (e.g., *Bx6*) is found in ancestral Panicoideae. The ancient Bx cluster was gained by the Triticeae ancestor via a horizontal transfer (HT) event from the ancestral Panicoideae and later separated into three parts on different chromosomes. *Bx6* appears to have been under less constrained selection during evolution of the Panicoideae as evidenced by the fact that was translocated ∼1.31-Mb away from the Bx cluster in *Z. mays*, moved to other chromosomes in *Echinochloa*, and even lost in *Dichanthelium*. Further investigation indicated that intense selection and polyploidization shaped the evolutionary trajectory of the Bx cluster in the grass family. This study provides the first case of HT of BGCs among plants and sheds new insights on the evolution of BGCs.

**Significance:** Biosynthetic gene clustering and horizontal gene transfer are two evolutionary inventions for rapid adaption by organisms. Horizontal transfer of a gene cluster has been reported in fungi and bacteria, but not in plants up to now. By mining the genomes of 40 monocot species, we deciphered the organization of Bx gene cluster, a biosynthetic gene cluster for benzoxazinoids in grasses. We found that the Bx cluster was formed by gene duplication of native analogues of individual Bx genes and directional translocation. More importantly, the Bx cluster in Triticeae was inherited from the Panicoideae via horizontal transfer. Compared with the native analogues, Bx clusters in grasses show constrained purifying selection underscoring their significance in environmental adaption.

## Introduction

Biosynthetic gene clusters (BGCs) are specialized genomic organizations comprised of a cluster of non-homologous genes contributing to the biosynthesis of chemical defensive metabolites (Nützmann et al., 2018; Nützmann and Osbourn, 2014). The selective advantages of clustering, such as gene co-regulation and co-inheritance, may promote the formation of BGCs (Rokas et al., 2018; Nützmann and Osbourn, 2014; Nützmann et al., 2016). Natural selection has also driven the establishment and maintenance of BGCs, including long-term purifying selection, positive selection, and balancing selection (Rokas et al., 2018; Slod and Rokas, 2010; Carbone et al., 2007; Liu et al., 2020; Takos and Rook, 2012). The formation and evolution of BGCs have been studied extensively in fungi (Rokas et al., 2018). About 30 examples of BGCs in plants have been identified in recent years (Guo e al., 2018). The Bx cluster for the biosynthesis of benzoxazinoids is the first identified BGC in plants (Frey et al., 1997).

Gene duplication, neofunctionalization and relocation have been suggested as the origins of BGCs in most fungi and plants (Nützmann et al., 2018; Rokas et al., 2018). The DAL gene cluster involved in the allantoin metabolism originated from duplication of native genes and relocation in the yeast *Saccharomyces cerevisiae* (Wong and Wolfe, 2005). The GAL cluster found in *Candida* yeasts originated through the relocation of native unclustered genes (Slot and Rokas, 2010). Horizontal transfer (HT) also leads to the emergence and spread of BGCs and is an important source of genomic innovation (Khaldi et al., 2008; Slot and Rokas, 2011; Reynolds et al., 2018; Kominek et al., 2019). In the fungus *Aspergillus clavatus*, the ACE1 gene cluster originated by HT from a donor closely related to the rice blast fungus *Magnaporthe grisea* (Khaldi et al., 2008). The GAL cluster of *Schizosaccharomyces* yeasts was acquired from a *Candida* yeast (Slot and Rokas, 2010). A full operon encoding siderophore biosynthesis genes was horizontally transferred from bacteria to a group of budding yeasts (Kominek et al., 2019). In animals, bdelloid rotifers, small freshwater invertebrates, appear to have acquired a BGC for cell wall peptidoglycan biosynthesis comprised of a racemase and a ligase, from bacteria (Gladyshev et al., 2008). In plants, BGCs were not likely to be derived from microbes via HT (Nützmann et al., 2018) and no BGCs via HT have been identified.

Benzoxazinoids are a class of indole-derived protective and allelopathic secondary metabolites that function in plants to defend against insect herbivores, microbial pathogens and neighboring competing plants (reviewed in Frey et al., 2009). 2,4-dihydroxy-1,4-benzoxazin-3-one (DIBOA) and its 7-methoxy analog DIMBOA are the predominant representatives of benzoxazinoids in plants (Frey et al., 1997; Frey et al., 2009) and these compounds have been identified in many plants, including maize (*Zea mays*), wheat (*Triticum aestivum*), and barnyardgrass (*Echinochloa crus-galli*) (Frey et al., 2009; Guo et al., 2017). In *Echinochloa*, a weed species, DIBOA functions as an allelopathic compound against rice in paddy fields (Guo et al., 2017).

The pathway of benzoxazinoid biosynthesis has been elucidated extensively in *Z. mays* (Fig. 1a). The first step is the biosynthesis of indole from indole-3-glycerolphosphate in the chloroplast by Bx1, a homolog of the α-subunit of tryptophan synthase. Four P450 monooxygenases from CYP71C subfamily (Bx2 to Bx5) add four oxygen atoms at four position of the indole to synthesize DIBOA, the simplest benzoxazinoid (Frey et al., 1997). Two UDP-glucosyltransferases (UGTs), Bx8 and Bx9, attach a glucose moiety to DIBOA to produce DIBOA-Glc (Rad et al., 2001). Bx6, a 2-oxoglutarate-dependent dioxygenase (2-ODD), oxidizes DIBOA-Glc to TRIBOA-Glc and subsequently Bx7 (OMT, O-methyltransferase) methylates TRIBOA-Glc to produce DIMBOA-Glc (Jonczyk et al., 2008). Four OMTs (Bx10 to Bx12, and Bx14) catalyze the conversion of DIMBOA-Glc to HDMBOA-Glc with functional redundancy (Meihls et al., 2013). Bx13, a Bx6-like 2-ODD, converts DIMBOA-Glc to TRIMBOA-Glc and TRIMBOA-Glc is further methylated to produce DIM2BOA-Glc by Bx7 (Handrick et al., 2016). Bx14 catalyzes the reaction from DIM2BOA-Glc to HDIM2BOA-Glc by methylation (Handrick et al., 2016).

**Fig. 1.**
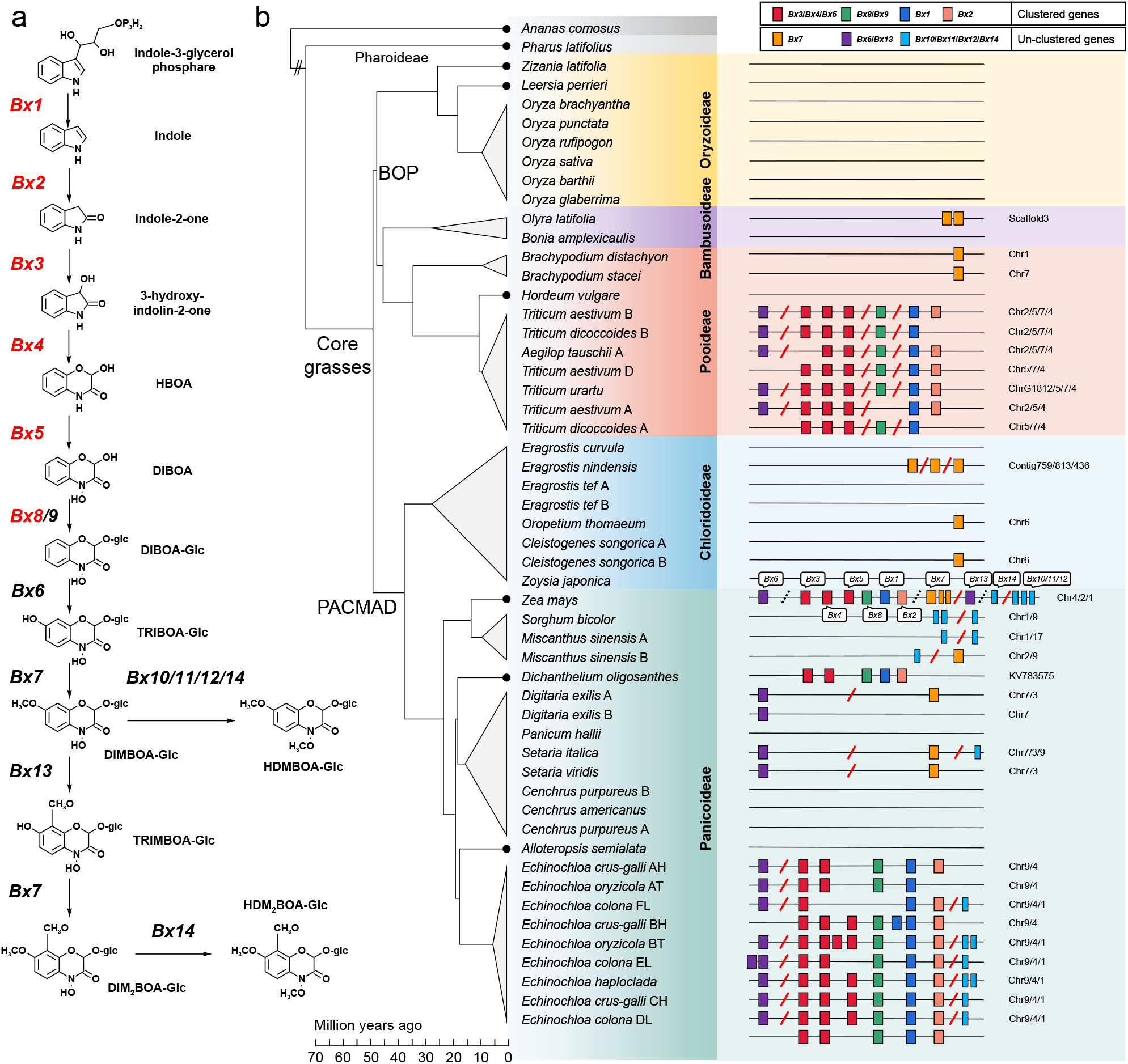
Benzoxazinoid biosynthesis pathway and distributions of Bx genes in grass. (a) Biosynthesis pathway of benzoxazinoid secondary metabolites in maize. The pathway(Bx)-related genes in the Bx cluster are marked in red. (b) Phylogeny and Bx gene distribution of grass species. Background colors represent different sub-families in Poaceae. The lineage divergence time is adopted from TimeTree database (www.timetree.org). Each rectangle represents one gene element. A red slash refers two different chromosomes for the neighbouring genes and a black dashed slash refers to a same chromosome but not clustered.

In maize, six Bx genes (*Bx1*-*Bx5*, and *Bx8*) encoding enzymes functioning in the first few steps of DIMBOA biosynthesis form a well-defined BGC (named the Bx cluster), located at the tip region of chromosome 4 (Frey et al., 1997; Frey et al., 2009). Bx genes have been identified in barnyardgrass within an intact cluster, and in wheat and rye within disperse sub-clusters (Guo et al., 2017; Sue et al., 2011). Previous studies indicated a monophyletic origin of Bx genes in benzoxazinoid biosynthesis (Frey et al., 2009; Sue et al., 2011; Dutartre et al., 2012; Nützmann and Osbourn, 2014). The progenitors have evolved Bx genes before the divergence of the Triticeae and the Panicoideae (Sue et al., 2011; Dutartre et al., 2012). However, it should be noted that limited sampling may bring over-interpretation of gene phylogeny. The frequent gene loss and rearrangements, and patchy distribution across divergent species complicated the understanding to the evolution of BGCs (Lind et al., 2017). The broad availability of high-quality genomes of important crops and wild grasses has facilitated the discovery of more BGCs (Guo et al., 2018) enabling us to trace the organization and evolution of BGCs more comprehensively and reliably.

Here, we identified all Bx genes in the grass family using 40 high-quality monocot genomes and further explored the origin of the Bx cluster and reconstructed its evolutionary trajectory. Through analysis of sequence similarities, phylogenies and genomic synteny, we provide evidence that the Bx clusters currently observed in grasses originated from a complex evolution processes that included HT. The HT event and further intense selection shaped the presence of the Bx cluster in the grass family.

## Results

### Identification and distribution of Bx genes in the grass family

Key genes in the benzoxazinoid biosynthesis pathway of *Z. mays*, including those in the Bx cluster (*Bx1* to *Bx5, Bx8*) and Bx genes dispersed in the genome (*Bx6, Bx7, Bx9* to *Bx14*) were used as baits to search the Bx genes in the genomes of 39 other species, covering five subfamilies of core grasses (Bambusoideae, Oryzoideae and Pooideae from BOP lineage, and Chloridoideae and Panicoideae from PACMAD lineage) and basal group of Poaceae (*Pharus latifolius*) (Fig. 1b; Table S1). All analogues of Bx genes were identified based on their sequence similarities, phylogeny, and genomic physical positions (Fig. 1b; Table S2).

In addition to the Bx clusters previously reported in *Z. mays* and *Echinochloa* (Frey, 1997; Guo et al., 2017), a Bx cluster was also found in *Dichanthelium oligosanthes*, Scribner’s rosette grass, a C3 panicoid grass (Fig. 1b). In the Triticeae, the Bx cluster was split into three sub-clusters located on three different chromosomes. In total, 12 clusters were found in six grass species, of which 10 were in *Echinochloa* genus, with one cluster in each monoploid genome (except one subgenome in *Echinochloa colona* with two copies). The Bx gene orders in clusters were entirely consistent among *Z. mays, D. oligosanthes* and *Echinochloa*, implying a single origin of the Bx clusters (Fig. 1b). Although the Bx cluster was split in the Triticeae, the order of *Bx3* to *Bx5* were same as the Bx cluster in the Panicoideae, which showed potential close relationship between Bx genes in BOP and PACMAD lineages. *Bx6* was distant 1.31-Mb away from Bx cluster in *Z. mays* genome, although both the gene and cluster were located on chromosome 4. *Bx6* was also identified in *Digitaria* and *Setaria* from Panicoideae. *Bx6* was located on chromosome 2 in Triticeae and chromosome 9 in *Echinochloa. Bx7* was an ancient gene, distributed in both BOP and PACMAD lineages, in spite of massive loss.

### Formation and HT of the Bx cluster (*Bx1* to *Bx5* and *Bx8*)

With *P. latifolius* from Pharoideae (N1 in Fig. 2a) serving as an outgroup, the gene tree of *Bx1* was divided into two lineages of BOP and PACMAD, in line with the species tree (Fig. 2a). *Bx1* genes formed a monoclade, composed by *Bx1* copies from previous identified species with Bx clusters. To distinguish other Bx homologs from *Bx1* copies, we called the other *Bx1* homologs as *Bx1* analogues. The *Bx1* analogues were native and extraordinarily conserved across the grass family and were in good synteny among genomes (Fig. 2b). The *Bx1* analogue AET5Gv21022100 (N0 in Fig. 2a) in *Aegilop tauschii* from Pooideae subfamily was syntenic to the *Bx1* analogue Pl3g34340 (N1) in *P. latifolius* from the basal lineage of Poaceae, as well as the *Bx1* analogues LOC_Os3g58300 (N2) in *Oryza sativa* from Oryzoideae, Et_4A_034058 (N5) in *Eragrostis tef* from Chloridoideae, Sevir.9G054600 (N6) in *Setaria viridis* from Paniceae, Panicoideae, and Zm00008a005484 (N8) in *Z. mays* from Andropogoneae, Panicoideae. Sequence alignments showed they were conserved with the domain of tryptophan synthase (Fig. 2c; Fig. S1). In contrast to the native *Bx1* analogues, the clade of *Bx1* copies, which is nested between native analogues of Chloridoideae and Panicoideae and sister to native copies of Panicoideae, is an extra lineage-specific copy duplicated in the ancestor of Panicoideae (Fig. 2a). To ensure the lineage-specific duplication event, local synteny of *Bx1* was scanned between *Z. mays* and other genomes (Fig. 2d). The two flanking genomic regions of Bx cluster in *Z. mays* showed high synteny to *Brachypodium distachyon* and *A. tauschii* from Pooideae, *O. sativa* from Oryzoideae, *Sorghum bicolor* from Andropogoneae, Panicoideae and *S. viridis* from Paniceae, Panicoideae. However, the Bx cluster was entirely absent in these genomes. Considering the species phylogeny in the Poaceae, the presence of Bx cluster in *Z. mays* is not ancestral but rather derived likely by translocation from other genomic positions. Comparing the gene positions of Bx clusters between *Z. mays* and *Echinochloa haploclada* from *Echinochloa*, their Bx clusters were in a large syntenic block, and the orders of Bx genes were consistent, implying the common origin of Bx cluster in their common ancestor before the divergence of Andropogoneae and Paniceae, although there was a translocation between them. Although the scaffold harboring the Bx cluster in *D. oligosanthes* was short, five Bx genes were assembled and their orders were in line with those in *Z. mays*, further supporting the origin of Bx cluster in ancestral Panicoideae. Sequence alignment among *Bx1* genes and their native analogues showed *Bx1* lineage-specific deletion and substitution, confirming a single origin of *Bx1* genes (Fig. 2c).

**Fig. 2.**
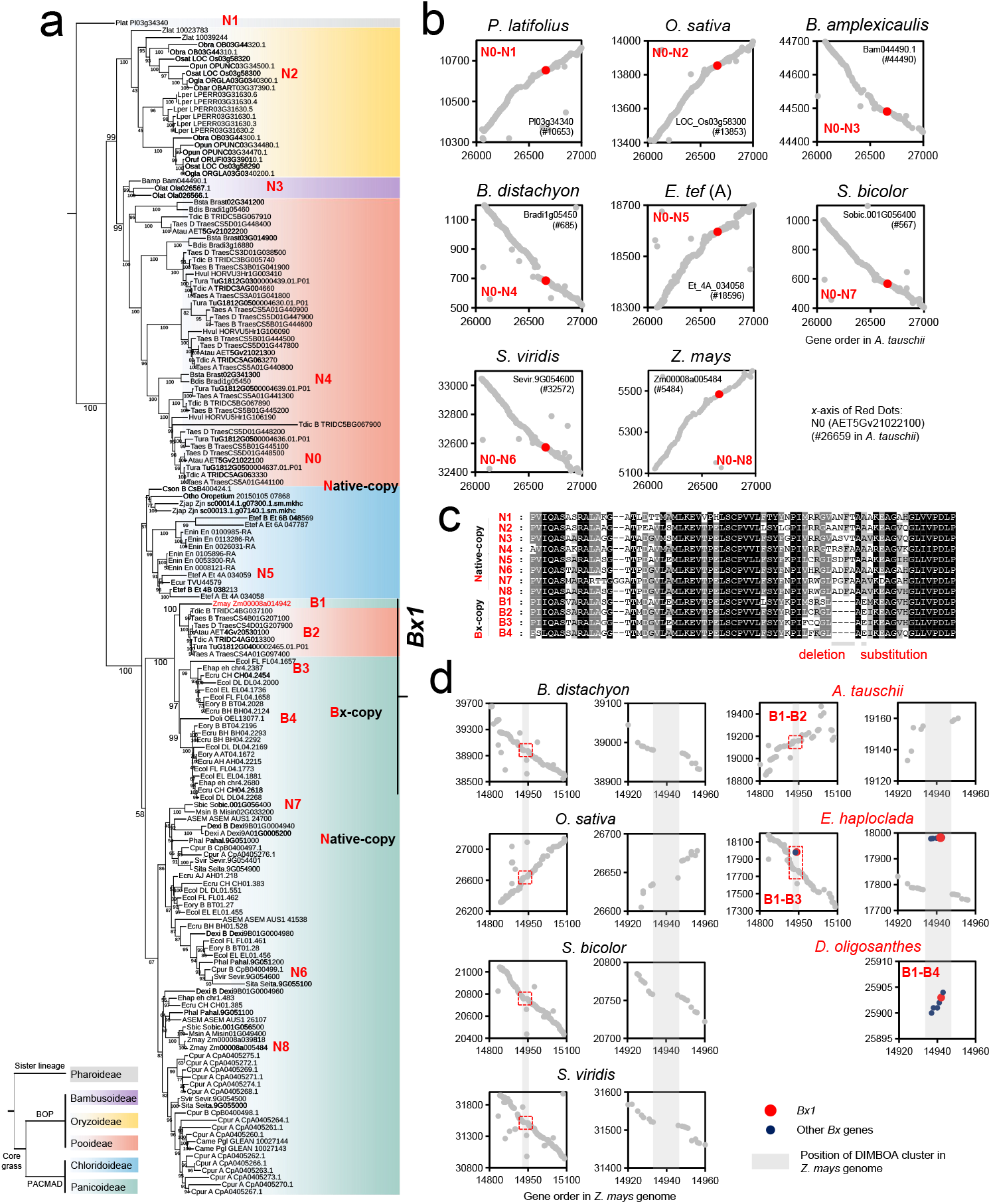
Phylogeny and genomic synteny of *Bx1* in grass. (a) Maximum-likelihood phylogenetic trees of *Bx1* in grass with *P. latifolius* as an outgroup species. Bootstrap value of 1000 replicates is labeled at each branch. The node label is composed of genome abbreviation and gene ID. Background filled colors represent subfamilies. The *Bx1* clade is highlighted as Bx-copy (e.g. B1-B4) and the paralogs of *Bx1* are labeled as native-copy (e.g. N0-N7). Left bottom tree shows the phylogenetic relationship of five subfamilies. (b) Genomic synteny among native *Bx1* analogues between species. Red dots represent the native *Bx1* analogues are syntenic. (c) Local protein sequence alignments among *Bx1* genes and their native analogues. Bx-copy specific deletion and amino acid substitution are marked in gray rectangles. (d) Genomic synteny between *Z. mays* and other species around the position of *Bx1*. For each species, the local synteny around *Bx1* is zoomed in at the right panel.

Within the *Bx1* clade, *Bx1* genes from the Triticeae form a monoclade nested among *Bx1* genes from Panicoideae, indicating a single origin of these genes. Given that the divergence between the Pooideae and Panicoideae is ancient, estimated at more than 50 million years ago (Ma et al., 2021) and native *Bx1* analogues are present, the positional congruence of the Triticeae *Bx1* clade is not likely to be derived from sexual hybridization, incomplete lineage sorting (ILS) or convergent evolution, but HT from the Panicoideae (Fig. 2a). To further confirm the robustness of the phylogeny of *Bx1* based on protein sequences, the phylogenetic trees of *Bx1* based on coding sequence (CDS), codon12 (first and second codon positions) and codon3 (third codon position) were built and the topologies confirmed the existence of gene duplication and HT of *Bx1* (Fig. S2).

We built the phylogeny and scanned the genomic synteny of *Bx2*-*Bx5* and *Bx8* across the whole Poaceae (Figs. S3 and S4). Native analogues of *Bx2* could be traced and were highly conservative (Fig. S3). *Bx3, Bx4* and *Bx5* were three tandemly duplicated CYP71C genes from cytochrome P450 superfamily. The native ancestral analogues of *Bx3* to *Bx5* were massively lost but the retained analogues showed high genomic synteny among subfamilies (Fig. S3). Based on the phylogeny of *Bx8, Bx8* genes were products of native analogues and *Bx8* genes in the Triticeae were nested within those in Panicoideae. *Bx9* was a maize-specific duplicate of *Bx8* (Fig. S4). In brief, topologies of the five Bx genes (*Bx2*-*Bx5* and *Bx8*) were similar to what was observed with *Bx1*, implying that Bx genes in the cluster were derived from a single origin and Bx genes in Triticeae were likely acquired via HT of an intact Bx cluster from Panicoideae.

To formally test the hypothesis of a Panicoideae origin of the Bx genes in Triticeae, we reconstructed phylogenies under constraints that the Bx genes in Triticeae were derived from Panicoideae Bx clade origin (PO) or outside of that clade (Non-PO). To determine whether the PO phylogenies statistically were better explanations than non-PO phylogenies we employed the approximately unbiased (AU) test, the resampling estimated log-likelihood method (RELL), and the Shimodaira-Hasegawa (SH) test. All tests of all Bx genes in cluster (*Bx1*-*Bx5* and *Bx8*) strongly rejected the alternative hypothesis that Bx genes in Triticeae were not derived from Panicoideae (all p values < 0.001 for AU tests) (Table S3). The results indicated that the obtained tree topologies of all Bx genes were highly robust and reflected a HT event of Bx genes from Panicoideae to Triticeae.

### Co-evolution between *Bx6* and the Bx cluster

The *Bx6* gene whose encoded product is responsible for oxidizing DIBOA-Glc to TRIBOA-Glc, the subsequent enzymatic step following the activity of the Bx cluster genes in maize was located away from the Bx cluster (Fig. 1a). The phylogeny of *Bx6* showed a similar pattern as *Bx1*, in that the Bx clade was duplicated from native *Bx6* analogues and HT from Panicoideae was likely responsible for the inheritance of the *Bx6* genes in Triticeae (Fig. 3a). Multi-species genome synteny analyses supported the above results (Fig. 3b). Topology tests confirmed the robustness of *Bx6* phylogeny and *Bx6* genes in Triticeae were nested within Panicoideae *Bx6* clade (p value < 0.001 for AU test) (Table S3). Hence, it is reasonable to speculate that *Bx6* co-evolved with the Bx cluster with similar evolutionary trajectories. Notably, besides species harboring Bx cluster, *Bx6* genes could be identified in *Setaria* and *Digitaria* from Panicoideae (Fig. 1b; Fig. 3a). The wide distribution of *Bx6* across the Panicoideae implies that *Bx6* originated by duplication at the common ancestor of Panicoideae. We also observed that *Bx13* is a maize-specific duplicate of *Bx6* (Fig. 3a).

**Fig. 3.**
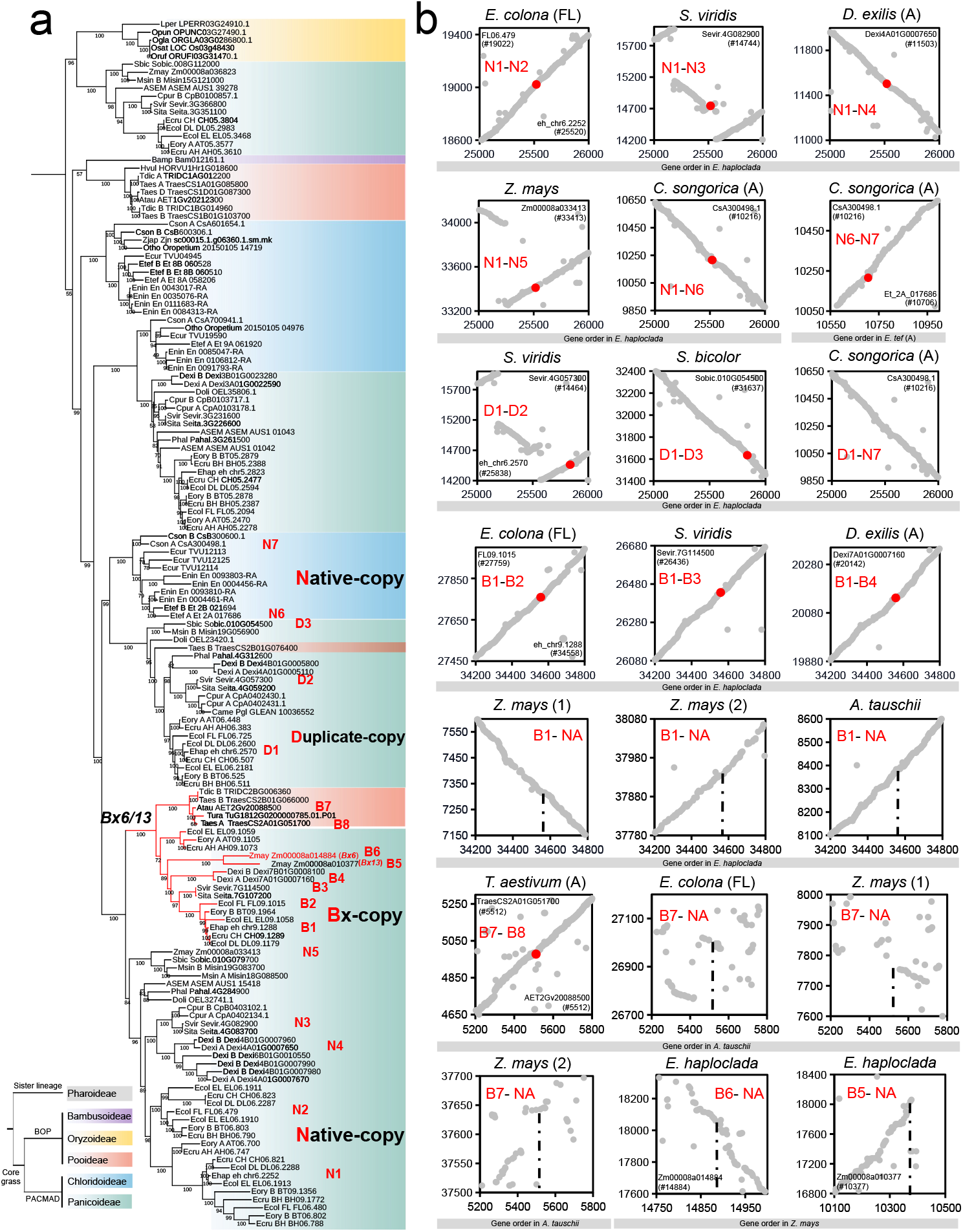
Phylogeny and genomic synteny of *Bx6* in grass. (a) Maximum-likelihood phylogenetic trees of *Bx6* in grass. Bootstrap value of 1000 replicates is labeled at each branch. Background filled colors represent subfamilies. The *Bx6* clade is highlighted as Bx-copy (e.g. B1-B8) and the native analogues of *Bx6* were labeled as native-copy (e.g. N1-N7). The other duplicates of *Bx6* native analogues are labeled as duplicate-copy (e.g. D1-D3). Left bottom tree shows the phylogenetic relationship of five subfamilies. (b) Genomic synteny among *Bx6* genes and their analogues between species based on gene order in each genome. Red dots represent the *Bx6* genes or analogues are syntenic in genome.

We also identified the presence of other dispersal Bx genes and built phylogenetic trees to trace their evolutionary histories. Bx7 catalyzes the conversion of TRIBOA-Glc to DIMBOA-Glc (Fig. 1a). Only limited homologs could be identified in grasses and its phylogenetic tree revealed that *Bx7* was conserved in evolution without congruence to species phylogeny, although massive losses occurred (Fig. S5). *Bx10*/*Bx11*/*Bx12*/*Bx14* encoded OMTs, acting as metabolic switches between caterpillar and aphid resistance, by transforming DIMBOA-Glc to HDMBOA-Glc (Li et al., 2018). From the phylogenetic analyses, the clade of *Bx10*/*Bx11*/*Bx12*/*Bx14* were maize-specific duplicates, and was in a well-defined Panicoideae-specific clade (Fig. S6). Within the clade, no Triticeae homologs were found. While in wheat (*T. aestivum*), two OMT genes were characterized as functional DIMBOA-Glc OMTs both designated as *TaBx10* but phylogenetically close to *Bx7*, rather than *Bx10* in *Z. mays*, indicating the convergence in function of OMT genes in grasses during the process of O-methylation (Li et al., 2018). This case implied that other paralogs of OMTs could function as *Bx10*/*Bx11*/*Bx12*/*Bx14* in the process of O-methylation and *Bx10*/*Bx11*/*Bx12*/*Bx14* are not compulsory for benzoxazinoid biosynthesis. Taken together, *Bx7* and *Bx10*/*Bx11*/*Bx12*/*Bx14* were alternative and dispensable to some extents in the Bx pathway. Hence in the following analyses, we focused on the Bx cluster and *Bx6*.

### Constrained purifying selection on the Bx cluster

Natural selection shapes the evolutionary dynamics of BGCs (Rokas et al., 2018; Slod and Rokas, 2010; Liu et al., 2020). The selection pressure was measured by *ω* (dN/dS, the ratio between non-synonymous sites substitution and synonymous sites substitution) in each lineages of the individual Bx genes. Generally, both the Bx genes and their native analogues were under purifying selection (*ω* < 1). Compared to outgroup lineage N-Chloridoideae (native Bx analogues in Chloridoideae), constrained purifying selections were detected in all of the native Bx genes for Panicoideae, with the exception of *Bx6*. The native analogues of *Bx6* in Panicoideae suffered relaxed selection with a higher *ω* value, relative to other Bx native genes. Compared to the native analogues, the *ω* values were lower for the Bx genes in Panicoideae (B-Panicoideae) in cluster, while no difference in selection was found for *Bx6*, which was not clustered together with Bx cluster. This selection bias in Panicoideae corresponded to the presence-and-absence (PAV) of Bx genes and their analogues (Fig. S7). The loss of native analogues of Bx genes in cluster was more frequent than those of Bx genes in cluster, which mirrored the relaxed selection, especially for *Bx2, Bx5* and *Bx8*. The presence of *Bx6* native analogues was highly conserved, with one copy within one single analyzed genome, corresponding to unbiased selection pressure compared to *Bx6* genes (Fig. S7). While in Triticeae, all of the Bx genes exhibited constrained selection, despite the conserved presence of Bx native analogues (Fig. 4a; Fig. S7). Although Bx genes in Triticeae were inferred to be gained from Panicoideae, stronger selection was detected in Triticeae Bx genes than those in Panicoideae, especially for *Bx1* and *Bx6*. To eliminate the effects by biases from species sampling and PAV of Bx genes or native analogues, the selection pressure was measured focusing on *Echinochloa* and Triticeae. The results further confirmed the selection profiling of Bx genes (Fig. S8).

**Fig. 4.**
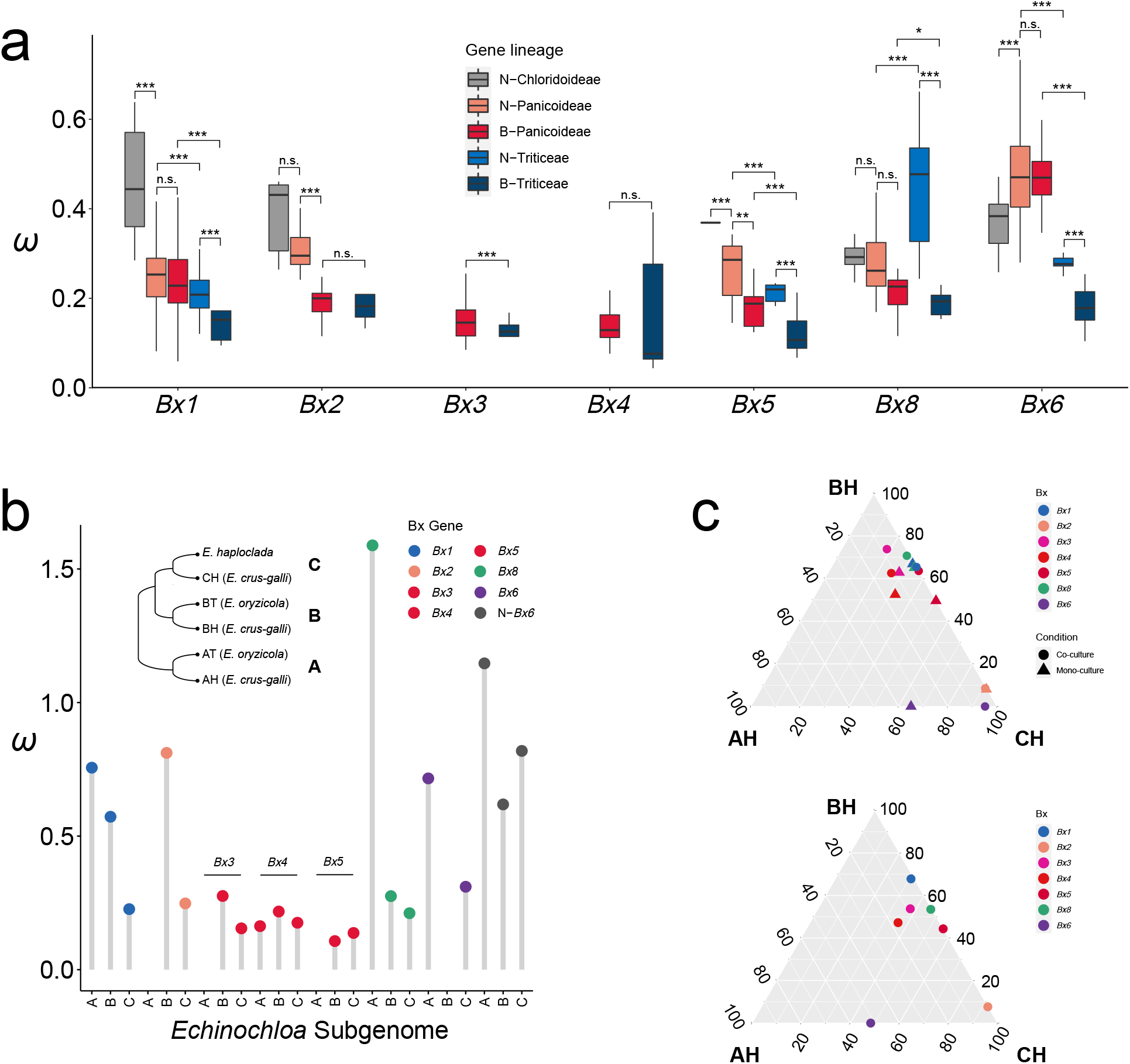
Selection and polyploidization effects on the Bx genes. (a) selection pressure estimated by *ω* of Bx genes and analogues. N-Chloridoideae, native analogues of Bx genes in Chloridoideae; N-Panicoideae, native analogues of Bx genes in Panicoideae; B-Panicoideae, Bx genes in Panicoideae; N-Triticeae, native analogues of Bx genes in Triticeae; B-Triticeae, Bx genes in Triticeae. In the box plots the horizontal line shows the median value, and the whiskers show the 25% and 75% quartile values of *ω*. Pairwise *t*-test are performed to evaluate significant differences. n.s., not significant; *, p < 0.05; **, p < 0.01; ***, p < 0.0001. (b) pairwise *ω* of Bx genes and analogues in subgenomes A, B and C between *E. crus-galli* and its progenitors (*E. haploclada* and *E. oryzicola*). The topology shows the phylogenetic relationship among subgenomes in the three *Echinochloa* species, where AT and AH belong to subgenome A, BT and BH belong to subgenome B, and *E. haploclada* and CH belonged to subgenome C. (c) relative expression (upper ternary diagram) and relative response contribution (lower ternary diagram) of multi-copy homologous Bx genes in *E. crus-galli* subgenomes (AH, BH and CH) under control and allelopathy treatment.

### Dominance of Bx cluster genes in polyploids

In species whose genomes contained Bx genes, polyploids are commonly seen (hexaploid *T. aestivum, E. crus-galli* and *E. colona*, and tetraploid *Triticum dicoccoides* and *Echinochloa oryzicola* in this study). We investigated the effects of polyploidization on Bx clusters or genes from three different views: PAV, selection, and gene expression. Duplicated genes tend to be lost due to gene redundancy or dosage effects in polyploids (Soltis and Soltis, 2009; Van de Peer et al., 2017). Not unexpectedly, Bx genes tended to be lost in polyploids, especially in *Echinochloa* (Fig.1b; Fig. S7). In diploid *Echinochloa haploclada*, the core Bx gene set was intact, while Bx losses were found in three polyploid *Echinochloa* species. In this case, only one intact copy of the core Bx gene set was retained in one subgenome in each species (e.g. BT in *E. oryzicola*, CH in *E. crus-galli* and DH2 in *E. colona*).

The selection strengths to homologous duplicates usually varies in polyploids (Ye et al., 2020). The genomes of *E. crus-galli* and its progenitors (*E. oryzicola* and *E. haploclada*) provided a model to study the selection dominance of multi-copy homologous Bx genes and we calculated the *ω* values of Bx genes in each subgenome between *E. crus-galli* and its parents (Fig. 4b). Bx genes in subgenome A were generally under relaxed purifying selection, with higher *ω* values compared to subgenomes B and C (e.g. *Bx1* and *Bx8*). For native analogues, the selection on subgenome A copy was relaxed in the example of *Bx6*. In general, biased selection was observed for Bx genes in *Echinochloa* and Bx genes in subgenome A were under less constrained selection in the post-hexaploidization.

Expression dominance has been commonly observed in polyploids (Ye et al., 2020; Van de Peer et al., 2017). Response contribution of subgenomes (relative changes of expressed transcripts from each subgenome, compared to the total expression change) is also biased among subgenomes (Ye et al., 2020). To explore the effect of polyploidization on gene expression of multi-copy Bx genes, we compared the expression levels of Bx genes in *E. crus-galli* with and without allelopathy treatment (i.e., co-culture with rice)(Guo et al., 2017). Expression and response contribution were both suppressed for Bx genes in subgenome AH (Fig. 4c). The dominance of selection and gene expression or response were associated such that Bx genes in subgenome A suffering less constrained selection, were suppressed in expression and response contribution (Fig. S8).

## Discussion

### Evolutionary trajectory of the Bx gene cluster in grass

Given that the Bx cluster and *Bx6* catalyze the first seven steps in the benzoxazinoid biosynthesis and are sufficient to synthesize benzoxazinoid compounds without other *Bx* genes (e.g. in wheat), we considered Bx cluster (*Bx1* to *Bx5* and *Bx8*) and *Bx6* as the core set of Bx genes in the pathway (Fig. 1a). Based upon the results from all of the phylogenetic analyses of core Bx genes, the evolutionary trajectory of Bx genes could be assumed (Fig. 5). Native Bx analogues could be found in all phylogenetic trees of core Bx genes and were evolutionarily conserved with good genomic synteny among subfamilies. Therefore, the Bx genes in the Bx cluster and *Bx6* should originate from duplication of native Bx analogues. Previous studies proposed *Bx1* evolved from duplication and modification of the alpha subunit of the tryptophan synthase (TSA) (Grun et al., 2005; Frey et al., 2009). Here, we comprehensively identified the native analogues of the Bx genes. Gene duplication, followed by neofunctionalization and/or subfunctionalization, and recurrent genomic translocation, gathered Bx genes together to form Bx cluster (Fig. 5). The processes of gene duplication and translocation may have been induced by activities of retrotransposon elements.

**Fig. 5.**
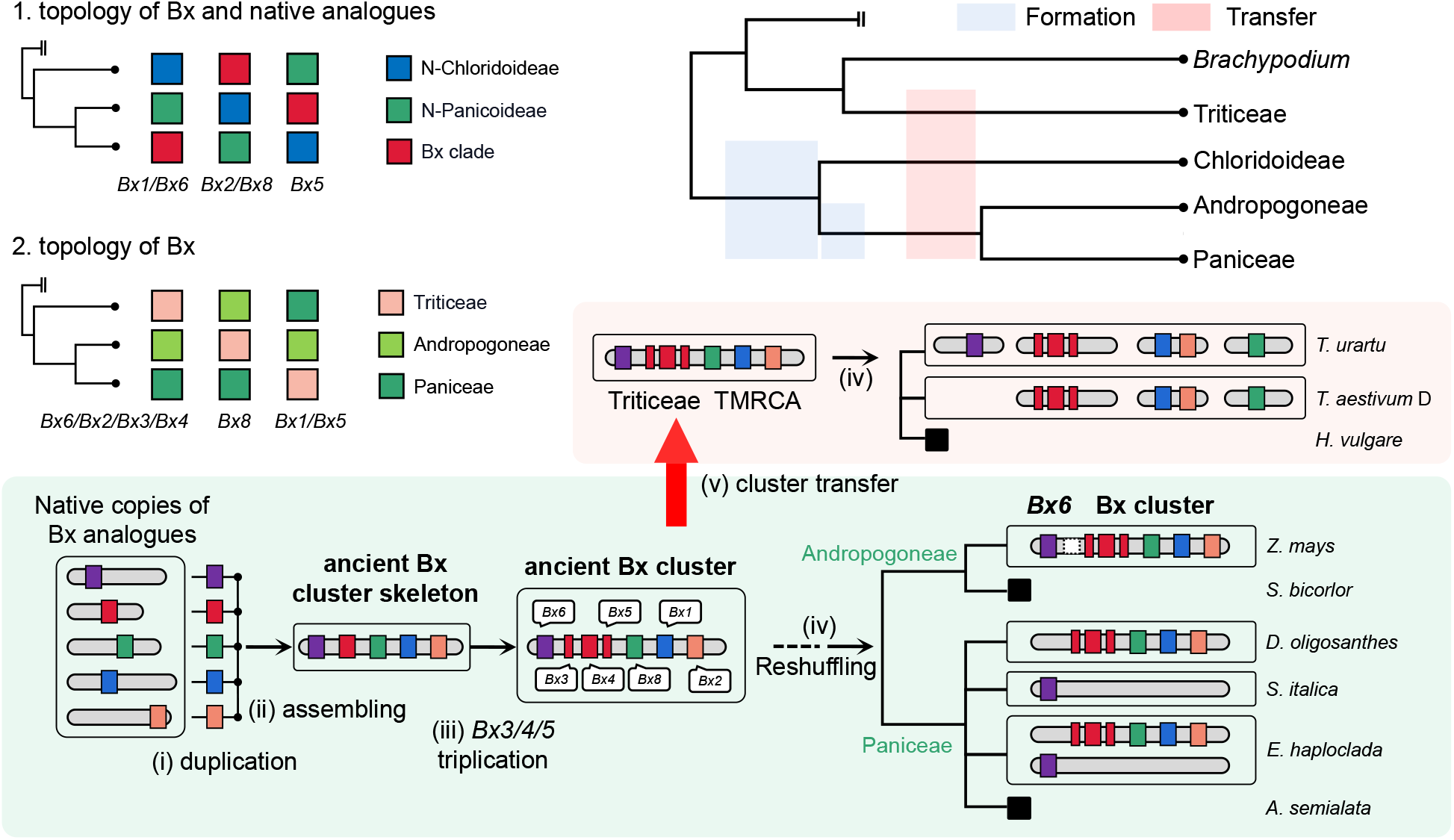
A proposed scenario for origin and evolution of the Bx cluster in grass. Top left shows different topologies of Bx genes or analogue in different lineages. Top right shows the relative divergence time of grass lineages. Blue shades represent the potential time range when Bx cluster was organized. Pink shade represents potential time range when the ancient Bx cluster (the current Bx cluster+*Bx6*) was transferred to Triticeae. Bottom shows the evolutionary trajectories of core Bx genes. TMRCA, The most recent common ancestor.

The positional relationship between Bx cluster and *Bx6* appears to be dynamic. *Bx6* and the Bx cluster are both located on chromosome 4 in *Z. mays* while they are separated into different chromosomes in Triticeae and *Echinochloa* (Fig. 1b). However, given that the genes in Bx cluster and *Bx6* showed almost the same evolutionary phylogenies and Bx6 catalyzes the reaction following those catalyzed by the gene products encodedin the Bx cluster, we speculated that *Bx6* co-evolved with the Bx cluster and were located in an ancient Bx cluster (Fig. 5). It is difficult to date accurately when the ancient Bx cluster formed, due to the unreliability of dating based on individual genes. However, we could infer that the duplication of Bx genes occurred at the common ancestor of Paniceae and Andropogoneae (supported by the *Bx1* and *Bx6* phylogenetic trees) or a common ancestor of Panicoideae and Chloridoideae (supported by the *Bx2* and *Bx8* phylogenies), and the Bx cluster was organized before the divergence of Paniceae and Andropogoneae.

Previously it was proposed that the genes of Bx biosynthesis in the grasses were of monophyletic origin before the divergence of the Triticeae and Panicoideae (Frey et al., 2009; Grun et al., 2005). Here, the integrated evidence indicates strongly that the Bx genes in Triticeae originated from Panicoideae via HT (Fig. 5). Triticeae and Panicoideae diverged more than 50 mya, which ruled out the possibility of natural hybridization between them and ILS. Previous studies also found that no benzoxazinoid biosynthesis can be detected in *Brachypodium* (basal genus in Pooideae) (Frey et al., 2009), corresponding to the absence of identifiable Bx genes in two *Brachypodium* genomes (Fig. 1b). Benzoxazinoids could be produced in wild *Hordeum* but not in cultivated *Hordeum* (*H. vulgare* in Triticeae), indicating the Bx genes were retained in wild *Hordeum* but lost in cultivated *Hordeum* (Grun et al., 2005; Sue et al., 2011). Therefore, it was speculated that the transfer occurred at the common ancestor of Triticeae after the divergence with *Brachypodium*. To trace the potential donor of Bx genes, we considered the topology between Bx genes of Triticeae, Andropogoneae (e.g. *Z. mays*) and Paniceae (e.g. *Echinochloa, D. oligosanthes*) (Fig. 5). Four Bx genes supported the common ancestor of Andropogoneae and Paniceae as the donor of Bx genes in Triticeae. However, three Bx genes showed discordant topology, implying the transfer event may have taken place at a time close to the divergence between Andropogoneae and Paniceae, which would result in an ILS-like phylogeny. With massive genome reshuffling in Triticeae, the intact ancient cluster (Bx cluster plus *Bx6*) was split into segments and scattered on four chromosomes (Frey et al., 2009). Gene loss resulted in the partial loss of Bx genes (e.g. *T. urartu*) and entire loss (e.g. *H. vulgare*) in Triticeae. It is noteworthy that phylogenies of individuals genes based on different sequence types (e.g., amino acid or nucleotide sequences), different substitution models, or other different parameters, are sometimes misleading. For example, the phylogenies of *Bx1* based on amino acid sequences and nucleotide sequences (CDS, codon12 and codon3) were incongruent since in Triticeae and Andropogoneae (*Z. mays*) *Bx1* formed a monoclade whereas *Bx1* formed a monoclade in Triticeae and Paniceae (Fig. 2a; Fig. S2).

In Panicoideae, genes in the Bx cluster and *Bx6* all showed a single common origin before the divergence of Andropogoneae and Paniceae based on data from analysis of gene phylogeny, genomic synteny and Bx gene orders. The common ancestor of Panicoideae had a cluster of Bx genes including *Bx6*. After the divergence between Andropogoneae and Paniceae, different genomic rearrangements happened in the two tribes (Fig. 5). In Andropogoneae, Bx cluster and *Bx6* were retained in *Z. mays* while being lost completely in other species (e.g., *S. bicolor* and *Miscanthus sinensis*). Furthermore, *Bx6* was separated away from the Bx cluster by translocation in *Z. mays*, although they were still on the short arm of chromosome 4. In Paniceae, massive losses were found in Bx genes. The Bx cluster was retained in *D. oligosanthes* but *Bx6* was lost. In contrast, *Bx6* was retained in *Setaria* and *Digitaria*, but *Bx* clusters were missing. Both Bx cluster and *Bx6* were absent in *Panicum, Cenchrus* and *Alloteropsis. Echinochloa* is the only genus in which the Bx cluster and *Bx6* are on two chromosomes (Fig. 1b).

### HT of gene cluster in plants

HT is an important driving force of trait innovation in various levels of organisms (Soucy et al., 2015). In plants, HT were commonly seen between parasites and corresponding host species, and between grafting rootstock and scion, due to intimate physical cell-to-cell contacts (Kim et al., 2014; Fuentes et al., 2014). HT could also emerge without direct contact, a phenomenon that has been studied somewhat in grasses (Hibdige et al., 2021; Dunning et al., 2019; Park et al., 2021). A total of 135 transferred candidate genes were identified across 17 grass species (Hibdige et al., 2021). Besides gene elements, transposon elements have also been detected to have been transferred among divergent grass species, as in the case for *Echinochloa* genus and *Oryza punctata* lineage (Park et al., 2021). In these reported HT events, a few have involved large genomic segments. A block containing 10 protein-coding genes was transmitted from *Iseilema membranaceum* (Andropopgoneae) to *Alloteropsis semialata* (Panicoideae) (Dunning et al., 2019). Here, we provided strong, unambiguous evidence that established that at least seven Bx biosynthetic genes in Triticeae are derived from donor ancestral Panicoideae as an intact ancient Bx cluster (including *Bx6*) via HT (Fig. 5). HT occurred more frequently between closely related species (Soucy et al., 2015; Hibdige et al., 2021), while Triticeae and Panicoideae were split more than 50 mya. The DNA transfer events from Panicoideae to Triticeae have been reported before. Several nuclear ribosomal DNA (rDNA) sequences in wild *Hordeum* and *Elymus* species were *Panicum*-like, indicating their foreign origins (Mahelka et al., 2010; Mahelka et al., 2017). Recently, a large chromosomal segment (∼68 kb long) harboring five stress-related protein-coding genes, has been reported to be transferred from *Panicum* to wild *Hordeum* species (Mahelka et al., 2021; Verhage, 2021). Some of these genes remained functional in the recipient *Hordeum* genomes. These cases reflected that the transfer of exotic DNA was not as rare among plants as previously supposed (Mahelka et al., 2021), at least in grass from Panicoideae to Triticeae. It is reasonable to infer that more HT events could be detected from Panicoideae to Triticeae in future studies and this unidirectional and biased HT pathway has accelerated the capacity to environmental stress in Triticeae.

Compared to prior reported plant-to-plant transfers, here we provide the first case of HT event of an intact gene cluster functioning in the biosynthesis of multi-effect chemical compounds in plants. The clustering of a series of biosynthetic genes facilitates the heritage and stress response by co-inheritance and co-expression in organisms, which is an ingenious invention in the long-term adaptive evolution. When combing HT and gene clustering together, it offers a rapid strategy to acquire highly efficient weapons to defend external stress. It seems this phenomenon is rare but universal in the kingdom of life, because transfers of BGC have been detected in fungi (Khaldi et al., 2008; Slot and Rokas, 2011; Reynolds et al., 2018). As for how the transfer between phylogenetically distant plant species occurs, one possible explanation is that it takes place because of occasional contact (e.g., like natural grafting) or is facilitated by vector transfer (e.g., insects, fungi, viruses) (Xia et al., 2021; Wang et al., 2020). The transfer of DNA between insect vectors and plants has been reported recently. For example, whitefly has acquired the plant-derived phenolic glucoside malonyltransferase gene *BtPMaT1* from a plant host enabling it to neutralize plant toxin phenolic glucosides (Xia et al., 2021). Similarly, the transfer of *Fhb7* from fungus *Epichloë* to *Thinopyrum* wheatgrass (Triticeae) provides broad resistance to both *Fusarium* head blight and crown rot in wheat (Wang et al., 2020).

### Selection on gene clusters

The driving forces for the organization and maintenance of BGCs remain in debate. Nevertheless, it is widely accepted that natural selection must inevitably shape their evolution. The selection analysis to BGCs were rare, due to limited identifications of BGCs and comparable sequences. In Saccharomycetes, the galactose BGCs are widely conserved in terms of sequence and function, suggesting the influence of long-term purifying selection (Slot and Rokas, 2010). Balancing selection also plays roles in maintaining diversity of BGCs, as in the case of the aflatoxin gene cluster in fungus *Aspergillus parasiticus* (Carbone et al., 2007). In *Arabidopsis*, the thalianol BGC appear to be under relaxed selection when compared with genes in the phytosterol biosynthetic pathway, but is still under strong purifying selection (Liu et al., 2020). In this study, we utilized multiple copies of Bx genes and their corresponding native analogues across a broad range of grass species to profile the selection landscapes of Bx clusters. Similar to what was found in the thalianol BGC, Bx genes in both Panicoideae and Triticeae showed purifying selection. When compared with native analogues, the selection on Bx genes in cluster was more constrained (Fig. 4a). The selection pressure was similar for *Bx6* and its native analogues in Panicoideae, possibly the result of dispersal of *Bx6* away from other core Bx genes in cluster. It is suggested that lateral pathway genes were less constrained than the early pathway genes in the biosynthesis of thalianol in *Arabidopsis* (Liu et al., 2020). Here, we noticed that *Bx6*, functioning after the reactions by genes in the Bx cluster, exhibited the highest *ω* value among the seven core Bx genes (Fig. 4a). *Bx8*, which is within Bx cluster, was less constrained than other Bx genes in the cluster. All identified Bx clusters or genes were transcribed in the various genomes and functioned in stress response, further indicating purifying selection in conserving the functions of Bx clusters.

### Subgenome dominance of gene clusters in polyploids

We found that several species identified to have Bx clusters or whole-set core Bx genes are polyploids (Fig. 1b). In most cases, polyploidization provides stronger growth and higher tolerance to environmental stress than original diploid status (Soltis and Soltis, 2009; Van de Peer et al., 2017). On this basis, biosynthetic gene clustering further offers these species a powerful weapon to response external stimulus. To some extent, the existence of BGCs in these polyploids assisted in allowing these species to become main crops under artificial selection (e.g., hexaploid and tetraploid wheat, and paleo-tetraploid maize) or successful agricultural weeds (hexaploid and tetraploid barnyardgrass). In polyploids, the subgenome dominance usually exists in selection and gene expression. The dominance of BGCs in polyploids has not been well studied. Differential expression of Bx genes in hexaploid wheat was detected (Nomura et al., 2005). The main contribution in hexaploid and tetraploid wheat is by subgenome B. In the hexaploid barnyardgrass *E. crus-galli*, we found an obvious suppression in expression of Bx genes on subgenome AH, compared with other two subgenomes (Fig. 4c). The dominance pattern of Bx genes was consistent with overall profiling across whole subgenomes with a significantly higher proportion of suppressed genes occurring in subgenome AH (Ye et al., 2020). Highly expressed metabolic genes tend to be retained preferentially after polyploidization due to selective pressure (Gout et al., 2009). The selection on Bx genes on subgenome A was indeed less constrained than that on other two Bx homologs (Fig. 4b; Fig. S8). Furthermore, three out of four Bx gene losses in the *E. crus-galli* pedigree were from subgenome A (Fig. S7). Gene loss is the extreme result of relaxed selection. Differential transposon element contents among three subgenomes may be one of the driving forces of expression suppression and relaxed selection on subgenome A in *Echinochloa*. More transposon elements on subgenome A somewhat increased the degree of methylation, which will inactivate the gene expression (Ye et al., 2020). As seen in the cases of wheat (Nomura et al., 2005) and barnyardgrass, the genomic bias in the expression of Bx genes in polyploids was putatively derived from the diploid progenitors. Subsequent selection would shape the presence-and-absence of Bx genes on each genome. Clearly, additional studies are needed to decipher the mechanism of dominance of BGCs in polyploids.

## Materials and Methods

### Datasets

Amino acid sequences of whole-genome protein and coding nucleotide sequences of 39 grass genomes (including grass basal group: *Pharus latifolius*; Oryzoideae: *Zizania latifolia, Leersia perrieri, Oryza brachyantha, Oryza punctata, Oryza rufipogon, Oryza sativa, Oryza barthii* and *Oryza glaberrima*; Bambusoideae: *Olyra latifolia* and *Bonia amplexicaulis*; Pooideae: *Brachypodium distachyon, Brachypodium stacei, Hordeum vulgare, Triticum aestivum, Triticum dicoccoides, Aegilop tauschii* and *Triticum urartu*; Chloridoideae: *Eragrostis curvula, Eragrostis nindensis, Eragrostis tef, Oropetium thomaeum, Cleistogenes songorica* and *Zoysia japonica*; Panicoideae: *Zea mays, Sorghum bicolor, Miscanthus sinensis, Dichanthelium oligosanthes, Digitaria exilis, Panicum hallii, Setaria italica, Setaria viridis, Cenchrus purpureus, Cenchrus americanus, Alloteropsis semialata, Echinochloa crus-galli, Echinochloa oryzicola, Echinochloa colona* and *Echinochloa haploclada*) and outgroup species *Ananas comosus* were downloaded from Phytozome (https://phytozome-next.jgi.doe.gov) and NGDC(https://ngdc.cncb.ac.cn)(Table S1). Polyploids with chromosome-level assemblies (hexaploid *T. aestivum, E. crus-galli* and *E. colona*, and tetraploid *T. dicoccoides, E. tef, C. songorica, M. sinensis, D. exilis, C. purpureus* and *E. oryzicola*) were split into subgenomes (Table S1). A total of 53 diploid or diploid-like genomes were used to construct grass phylogeny. OrthoFinder was used to identify single-copy orthologs in the 40 species genomes (Emms and Kelly, 2019). Individual phylogenetic trees of 45 single-copy genes were constructed using IQ-TREE (v1.6.12) with the best substitution model Model Finder (Nguyen et al., 2015) and integrated into a species tree using ASTRAL (v5.7.4) (Zhang et al., 2018). The divergence time was adopted from TimeTree database (www.timetree.org) (Kumar et al., 2017).

### Identification of Bx genes in grass

The protein sequences of Bx genes in *Z. mays* (*Bx1*, Zm00008a014942; *Bx2*, Zm00008a014943; *Bx3*, Zm00008a014937; *Bx4*, Zm00008a014938; *Bx5*, Zm00008a014940; *Bx6*, Zm00008a014884; *Bx7*, Zm00008a015292; *Bx8*, Zm00008a014941; *Bx9*, Zm00008a003056; *Bx10*, Zm00008a001636; *Bx11*, Zm00008a001638; *Bx12*, Zm00008a001639; *Bx13*, Zm00008a010377; *Bx14*, Zm00008a008314) were used as baits to search Bx genes in grass species by BLASTP. The homologs of individual Bx genes were filtered by parameters of *e*-value less than 1e-30 and identity greater than 50%. Homologs were then aligned using MAFFT (v7.310) (Katoh and Standley, 2013) and phylogenetic trees were built using IQ-TREE under the substitution model parameter ModelFinder with 1000 times of bootstrap replicates (Nguyen et al., 2015). Using the homologs in *A. comosus* or *P. latifolius* as outgroup, we only kept the closest homologous copies of Bx genes across the grass family as native analogues. For Bx trees where Bx homologs could not be found in outgroup species *A. comosus* and *P. latifolius*, we referred to the topological relationship among homologs in five subfamilies, to determine Bx genes and their native analogues.

### Phylogenetic analysis

Bx homologs (Bx genes and native analogues) were re-aligned using MAFFT (Katoh and Standley, 2013). Substitution models were selected using ModelFinder and the maximum-likelihood phylogenetic trees were reconstructed by IQ-TREE using Ultrafast Bootstrap Approximation (1000 replicates) for branch support (Nguyen et al., 2015). Tests of tree topologies, including RELL approximation, Shimodaira-Hasegawa (SH) test and approximately unbiased (AU) test, were performed using IQ-TREE with 10000 bootstrap replicates (Nguyen et al., 2015). To eliminate the effects of protein sequence alignment gaps, we also used Gblocks (Castresana, 2000) to remove gaps from alignments with parameter “-b4=5 -b5=h”. The trimmed alignments of conserved regions were used in topology tests. The phylogeny constructions of Bx genes based on coding sequence, codon12 (first and second positions within a codon) and codon3 (third position within a codon) were performed using MAFFT for alignment and IQ-TREE with the best substitution model (ModelFinder) and 1000-replicate ultrafast bootstrap analysis (Nguyen et al., 2015).

### Genome synteny analysis

Whole-genome protein sequences were compared pairwise among the 39 grass species using BLASTP. The best hit of each blast was kept. We also required that the *e*-value should be less than 1e-30 and identity greater than 50%. According to the physical positions of the genes on each chromosome of each species, the genes or proteins were ordered. We performed the gene-to-gene synteny analysis among grass species based on their orders within each genome.

### Selection analysis

Selection pressure was measured by indicator *ω*, the ratio between non-synonymous substitution rate (dN) and synonymous substitution rate (dS), with usually *ω* = 1 meaning neutral mutations, *ω* < 1 purifying selection, and *ω* > 1 diversifying positive selection. Bx homologs whose lengths of CDS or protein sequences were longer than two-times or shorter than half of the lengths of Bx genes or proteins in *Z. mays* were removed. Within each clade in the phylogenetic tree of each Bx gene, only one copy was kept in following analysis within one (sub)genome for tandem duplicates and the copy of duplicates with abnormal sequence length (usually much shorter) was removed. The CDS and protein sequences were aligned using MAFFT and PAL2NAL (Suyama et al., 2006). The dN and dS values were calculated using KaKs_calculator in the NG model for all pairs of genes within each clade (Bx clade or native analogue clade) (Zhang et al., 2006).

### Gene expression analysis

RNA-seq data from an analysis of *E. crus-galli* seedlings under the conditions of mono-culture and co-culture with rice were downloaded from NCBI (BioProject PRJNA268892) (Guo et al., 2017) and the low-quality reads were removed using the NGSQC toolkit (v2.3.348) (Patel and Jain, 2012). The clean reads were mapped to the chromosome-level reference genome of *E. crus-galli* (STB08) using TopHat (v2.1.1) (Trapnell et al., 2012). Relative gene expression levels were quantified and normalized to FPKM values using Cufflinks (v2.2.1) (Trapnell et al., 2012). The determination of expression dominance and response contributions of Bx genes in subgenomes of *E. crus-galli* followed a previously described approach (Ye et al., 2020).

## Acknowledgement

The authors are grateful to Jie Qiu for insightful suggestions on the manuscript. This work was supported by grants from the Zhejiang Natural Science Foundation (LZ17C130001), Jiangsu Collaborative Innovation Center for Modern Crop Production and 111 Project (B17039).

## Author contributions

L. F. and D.W. conceived and designed research. L.F. and C.-Y.Y. supervised the research. D.W. and B.J. carried out the data analysis. D.W., L.F., M.P.T, and C.-Y.Y. analyzed the findings and wrote the manuscript.

## Additional information

Supporting information is available for this paper online.

## Supporting information

Table S1. A list of plant genomes used in this study

Table S2. Core Bx genes (*Bx1* to *Bx6* and *Bx8*) and corresponding native analogues in grass genomes

Table S3. Topology tests of two hypothesis on transfer of Bx genes in Triticeae from Panicoideae

Fig. S1. Alignment of amino acid sequences of *Bx1* gene in Fig. 2c

Fig. S2. Phylogenies of *Bx1* based on CDS, codon12 and codon3 datasets, related to Fig. 2a.

Fig. S3. Phylogeny of *Bx2* to *Bx5* and genomic synteny of *Bx2* and *Bx5* regions across the grass family

Fig. S4. Phylogeny of *Bx8* and *Bx9* across the grass family

Fig. S5. Phylogeny of *Bx7* across the grass family

Fig. S6. Phylogeny of *Bx10* to *Bx12* and *Bx14* across the grass family

Fig. S7. Presence and absence of Bx genes (B-copy) and native analogues (N-copy). Blue grids represent presence and white represent absence.

Fig. S8. Selection pressure of Bx genes in Echinochloa and Triticeae. In the box plots the horizontal line shows the median value, and the whiskers show the 25% and 75% quartile values of *ω*. B-copy, Bx genes; N-copy, native analogues of Bx genes.

Fig. S9. Negative relationship between selection indicator *ω* values and expression or response dominance.

