## Supplementary material for "Horizontal Transfer and Evolution of the Biosynthetic Gene Cluster for Benzoxazinoid"

### Supplementary Tables and Figures

**Table S1. A list of plant genomes used in this study**

| Lineage | Sub-family | Species | Ploid | Assembly level | Version | Genome Abbr. |
|---|---|---|---|---|---|---|
| Outgroup | Bromeliaceae | *Ananas comosus* | 2 | chromosome | F153 | Acom |
| Pharoideae | Pharoideae | *Pharus latifolius* | 2 | chromosome | 60161 | Plat |
| BOP | Bambusoideae | *Olyra latifolia* | 2 | scaffold | / | Olat |
| | | *Bonia amplexicaulis* | 6 | scaffold | / | Bamp |
| | Oryzoideae | *Leersia perrieri* | 2 | chromosome | GCA_000325765.3 | Lper |
| | | *Oryza barthii* | 2 | chromosome | GCA_000182155.2 | Obar |
| | | *Oryza brachyantha* | 2 | chromosome | GCA_000231095.2 | Obra |
| | | *Oryza glaberrima* | 2 | chromosome | GCA_000576495.1 | Ogla |
| | | *Oryza punctata* | 2 | chromosome | GCA_000573905.1 | Opun |
| | | *Oryza rufipogon* | 2 | chromosome | GCA_000817225.1 | Oruf |
| | | *Oryza sativa* | 2 | chromosome | v7.0 | Osat |
| | | *Zizania latifolia* | 2 | scaffold | v01 | Zlat |
| | Pooideae | *Aegilop tauschii* | 2 | chromosome | v4.0 | Atau |
| | | *Brachypodium distachyon* | 2 | chromosome | v3.1 | Bdis |
| | | *Brachypodium stacei* | 2 | chromosome | v1.1 | Bsta |
| | | *Hordeum vulgare* | 2 | chromosome | / | Hvul |
| | | *Triticum aestivum* | 6 | chromosome | iwgsc_refseqv1.0 | Taes_A; Taes_B; Taes_D |
| | | *Triticum dicoccoides* | 4 | chromosome | WEWSeq_v.1.0 | Tdic_A; Tdic_B |
| | | *Triticum urartu* | 2 | chromosome | Tu | Tura |
| PACMAD | Panicoideae | *Alloteropsis semialata* | 2 | scaffold | AUS1_V1.0 | Asem |
| | | *Cenchrus americanus* | 2 | chromosome | Camericanus.prot | Came |
| | | *Cenchrus purpureus* | 4 | chromosome | GWHAORA00000000 | Cpur_A; Cpur_B |
| | | *Dichanthelium oligosanthes* | 2 | scaffold | / | Doli |
| | | *Digitaria exilis* | 4 | chromosome | CM05836 | Dexi_A; Dexi_B |
| | | *Miscanthus sinensis* | 4 | chromosome | v7.1 | Msin_A; Msin_B |
| | | *Panicum hallii* | 2 | chromosome | v3.1 | Phal |
| | | *Setaria italica* | 2 | chromosome | v2.2 | Sita |
| | | *Setaria viridis* | 2 | chromosome | v2.1 | Svir |
| | | *Sorghum bicolor* | 2 | chromosome | v3.1.1 | Sbic |

| | | | | | | |
|---|---|---|---|---|---|---|
| | | *Zea mays* | 2 | chromosome | PH207_v1.1 | Zmay |
| | | *Echinochloa haploclada* | 2 | chromosome | v1 | Ehap |
| | | *Echinochloa oryzicola* | 4 | chromosome | v2 | Eory_AT; Eory_BT |
| | | *Echinochloa crus-galli* | 6 | chromosome | v3 | Ecru_AH; Ecru_BH; Ecru_CH |
| | | *Echibochloa colona* | 6 | chromosome | v1 | Ecol_DL; Ecol_EL; Ecol_FL |
| | Chloridoideae | *Cleistogenes songorica* | 4 | chromosome | GWHANUQ00000000 | Cson_A; Cson_B |
| | | *Eragrostis curvula* | 2 | scaffold | / | Ecur |
| | | *Eragrostis tef* | 4 | chromosome | v3.1 | Etef_A; Etef_B |
| | | *Eragrostis nindensis* | 4 | contig | v2.1 | Enin |
| | | *Oropetium thomaeum* | 2 | contig | Othomaeum | Otho |
| | | *Zoysia japonica* | 4 | scaffold | Zjaponica_r1.1 | Zjap |

**Table S2. Core Bx genes (*Bx1* to *Bx6* and *Bx8*) and corresponding native analogues in grass genomes**

| Species | Genome | Bx | Bx copy | Chromosome | Native analogues |
|---|---|---|---|---|---|
| *A. tauschii* | | *Bx1* | AET4Gv20530100 | 4D | AET5Gv21022100 |
| | | | | | AET5Gv21022200 |
| | | | | | AET5Gv21021300 |
| *T. urartu* | | *Bx1* | TuG1812G0400002465.01 | Tu4 | TuG1812G0500004630.01 |
| | | | | | TuG1812G0500004636.01 |
| | | | | | TuG1812G0500004637.01 |
| | | | | | TuG1812G0500004639.01 |
| *T. dicoccoides* | A | *Bx1* | TRIDC4AG013300 | 4A | TRIDC5AG063330 |
| | | | | | TRIDC5AG063270 |
| *T. dicoccoides* | B | *Bx1* | TRIDC4BG037100 | 4B | TRIDC5BG067890 |
| | | | | | TRIDC5BG067900 |
| | | | | | TRIDC5BG067910 |
| *T. aestivum* | A | *Bx1* | TraesCS4A01G097400 | chr4A | TraesCS5A01G441100 |
| | | | | | TraesCS5A01G441300 |
| | | | | | TraesCS5A01G440800 |
| | | | | | TraesCS5A01G440900 |
| *T. aestivum* | B | *Bx1* | TraesCS4B01G207100 | chr4B | TraesCS5B01G445100 |
| | | | | | TraesCS5B01G445200 |
| | | | | | TraesCS5B01G444500 |
| | | | | | TraesCS5B01G444600 |
| *T. aestivum* | D | *Bx1* | TraesCS4D01G207900 | chr4D | TraesCS5D01G448400 |
| | | | | | TraesCS5D01G448500 |
| | | | | | TraesCS5D01G448200 |
| | | | | | TraesCS5D01G447800 |
| | | | | | TraesCS5D01G447900 |
| *Z. mays* | | *Bx1* | Zm00008a014942 | chr04 | Zm00008a005484 |
| *D. oligosanthes* | | *Bx1* | OEL13077.1 | KV783575.1 | |
| *E. haploclada* | | *Bx1* | eh_chr4.2680 | eh_chr4 | eh_chr1.483 |
| *E. oryzicola* | AT | *Bx1* | AT04.1672 | AT04 | |
| *E. oryzicola* | BT | *Bx1* | BT04.2196 | BT04 | BT01.27 |
| | | | | | BT01.28 |
| *E. crus-galli* | AH | *Bx1* | AH04.2215 | AH04 | AH01.218 |
| *E. crus-galli* | BH | *Bx1* | BH04.2292 | BH04 | BH01.528 |
| | | | BH04.2293 | | |
| *E. crus-galli* | CH | *Bx1* | CH04.2618 | CH04 | CH01.385 |
| | | | | | CH01.383 |
| *E. colona* | DL1 | *Bx1* | DL04.2268 | DL04 | DL01.551 |
| *E. colona* | DL2 | *Bx1* | DL04.2169 | DL04 | DL01.551 |
| *E. colona* | EL | *Bx1* | EL04.1881 | EL04 | EL01.455 |
| | | | | | EL01.456 |
| *E. colona* | FL | *Bx1* | FL04.1773 | FL04 | FL01.461 |
| | | | | | FL01.462 |
| *A. tauschii* | | *Bx2* | AET4Gv20530000 | 4D | |
| *T. urartu* | | *Bx2* | TuG1812G0400002466.01 | Tu4 | |

| | | | | | |
|---|---|---|---|---|---|
| *T. aestivum* | A | *Bx2* | TraesCS4A01G097500 | chr4A | |
| *T. aestivum* | B | *Bx2* | TraesCS4B01G207000 | chr4B | |
| *T. aestivum* | D | *Bx2* | TraesCS4D01G207800 | chr4D | |
| *Z. mays* | | *Bx2* | Zm00008a014943 | chr04 | |
| *D. oligosanthes* | | *Bx2* | OEL13078.1 | KV783575.1 | OEL15381.1 |
| *E. haploclada* | | *Bx2* | eh_chr4.2681 | eh_chr4 | |
| *E. oryzicola* | AT | *Bx2* | | | AT08.48 |
| | | | | | AT08.49 |
| *E. oryzicola* | BT | *Bx2* | BT04.2195 | BT04 | BT08.44 |
| *E. crus-galli* | AH | *Bx2* | AH04.2216 | AH04 | |
| *E. crus-galli* | BH | *Bx2* | BH04.2291 | BH04 | |
| *E. crus-galli* | CH | *Bx2* | CH04.2619 | CH04 | |
| *E. colona* | DL1 | *Bx2* | DL04.2269 | DL04 | |
| *E. colona* | DL2 | *Bx2* | DL04.2172 | DL04 | |
| *E. colona* | EL | *Bx2* | EL04.1882 | EL04 | |
| *E. colona* | FL | *Bx2* | FL04.1774 | FL04 | |
| *T. urartu* | | *Bx3* | TuG1812G0500000122.01 | Tu5 | |
| *T. dicoccoides* | A | *Bx3* | TRIDC5AG001550 | 5A | |
| *T. dicoccoides* | B | *Bx3* | TRIDC5BG001410 | 5B | |
| *T. aestivum* | A | *Bx3* | TraesCS5A01G008900 | chr5A | |
| *T. aestivum* | B | *Bx3* | TraesCS5B01G007200 | chr5B | |
| *T. aestivum* | D | *Bx3* | TraesCS5D01G014300 | chr5D | |
| *Z. mays* | | *Bx3* | Zm00008a014937 | chr04 | |
| *D. oligosanthes* | | *Bx3* | OEL13081.1 | KV783575.1 | |
| *E. haploclada* | | *Bx3* | eh_chr4.2676 | eh_chr4 | |
| *E. oryzicola* | AT | *Bx3* | AT04.1669 | AT04 | |
| *E. oryzicola* | BT | *Bx3* | BT04.2201 | BT04 | |
| *E. crus-galli* | AH | *Bx3* | AH04.2211 | AH04 | |
| *E. crus-galli* | BH | *Bx3* | BH04.2297 | BH04 | |
| *E. crus-galli* | CH | *Bx3* | CH04.2613 | CH04 | |
| *E. colona* | DL1 | *Bx3* | DL04.2265 | DL04 | |
| *E. colona* | DL2 | *Bx3* | DL04.2165 | DL04 | |
| *E. colona* | EL | *Bx3* | EL04.1877 | EL04 | |
| *E. colona* | FL | *Bx3* | FL04.1772 | FL04 | |
| *A. tauschii* | | *Bx4* | AET5Gv20025200 | 5D | |
| *T. urartu* | | *Bx4* | TuG1812G0500000121.01 | Tu5 | |
| *T. dicoccoides* | A | *Bx4* | TRIDC5AG001540 | 5A | |
| *T. dicoccoides* | B | *Bx4* | TRIDC5BG001400 | 5B | |
| *T. aestivum* | A | *Bx4* | TraesCS5A01G008800 | chr5A | |
| *T. aestivum* | B | *Bx4* | TraesCS5B01G007100 | chr5B | |
| *T. aestivum* | D | *Bx4* | TraesCS5D01G014200 | chr5D | |
| *Z. mays* | | *Bx4* | Zm00008a014938 | chr04 | |
| *D. oligosanthes* | | *Bx4* | OEL13080.1 | KV783575.1 | |
| *E. haploclada* | | *Bx4* | eh_chr4.2677 | eh_chr4 | |
| *E. oryzicola* | AT | *Bx4* | AT04.1670 | AT04 | |
| *E. oryzicola* | BT | *Bx4* | BT04.2199 | BT04 | |
| | | | BT04.2200 | | |

| | | | | | |
|---|---|---|---|---|---|
| *E. crus-galli* | AH | *Bx4* | AH04.2212 | AH04 | |
| *E. crus-galli* | BH | *Bx4* | BH04.2296 | BH04 | |
| *E. crus-galli* | CH | *Bx4* | CH04.2614 | CH04 | |
| *E. colona* | DL1 | *Bx4* | DL04.2266 | DL04 | |
| *E. colona* | DL2 | *Bx4* | DL04.2166 | DL04 | |
| *E. colona* | EL | *Bx4* | EL04.1878 | EL04 | |
| *A. tauschii* | | *Bx5* | AET5Gv20025100 | 5D | AET2Gv21022600 |
| *T. urartu* | | *Bx5* | TuG1812G0500000120.01 | Tu5 | TuG1812G0200005151.01 |
| *T. dicoccoides* | A | *Bx5* | TRIDC5AG001530 | 5A | TRIDC2AG065830 |
| *T. dicoccoides* | B | *Bx5* | TRIDC5BG001390 | 5B | |
| *T. aestivum* | A | *Bx5* | TraesCS5A01G008700 | chr5A | |
| *T. aestivum* | B | *Bx5* | TraesCS5B01G007000 | chr5B | TraesCS2B01G485100 |
| *T. aestivum* | D | *Bx5* | TraesCS5D01G014100 | chr5D | TraesCS2D01G464200 |
| *Z. mays* | | *Bx5* | Zm00008a014940 | chr04 | |
| *E. haploclada* | | *Bx5* | eh_chr4.2678 | eh_chr4 | eh_chr8.27 |
| *E. oryzicola* | BT | *Bx5* | BT04.2198 | BT04 | |
| *E. crus-galli* | BH | *Bx5* | BH04.2295 | BH04 | |
| *E. crus-galli* | CH | *Bx5* | CH04.2615 | CH04 | |
| *E. colona* | DL2 | *Bx5* | DL04.2167 | DL04 | |
| *E. colona* | FL | *Bx5* | | | FL02.2320 |
| *A. tauschii* | | *Bx8* | AET7Gv20307400 | 7D | AET7Gv20836200 |
| *T. urartu* | | *Bx8* | TuG1812G0700001239.01 | Tu7 | TuG1812G0700003832.01 |
| *T. dicoccoides* | A | *Bx8* | TRIDC7AG014040 | 7A | |
| *T. dicoccoides* | B | *Bx8* | TRIDC1BG074840 | 1B | |
| *T. aestivum* | A | *Bx8* | | | TraesCS7A01G344300 |
| *T. aestivum* | B | *Bx8* | TraesCS7B01G016800 | chr7B | TraesCS7B01G238800 |
| *T. aestivum* | D | *Bx8* | TraesCS7D01G116700 | chr7D | TraesCS7D01G335200 |
| *Z. mays* | | *Bx8* | Zm00008a014941 | chr04 | |
| *D. oligosanthes* | | *Bx8* | OEL13079.1 | KV783575.1 | |
| *E. haploclada* | | *Bx8* | eh_chr4.2679 | eh_chr4 | |
| *E. oryzicola* | AT | *Bx8* | AT04.1671 | AT04 | |
| *E. oryzicola* | BT | *Bx8* | BT04.2197 | BT04 | |
| *E. crus-galli* | AH | *Bx8* | AH04.2214 | AH04 | |
| *E. crus-galli* | BH | *Bx8* | BH04.2294 | BH04 | |
| *E. crus-galli* | CH | *Bx8* | CH04.2616 | CH04 | |
| *E. colona* | DL1 | *Bx8* | DL04.2267 | DL04 | |
| *E. colona* | DL2 | *Bx8* | DL04.2168 | DL04 | |
| *E. colona* | EL | *Bx8* | EL04.1879 | EL04 | |
| *A. tauschii* | | *Bx6* | AET2Gv20088500 | 2D | AET1Gv20212300 |
| *T. urartu* | | *Bx6* | TuG1812G0200000785.01 | Tu2 | |
| *T. dicoccoides* | A | *Bx6* | | | TRIDC1AG012200 |
| *T. dicoccoides* | B | *Bx6* | TRIDC2BG006360 | 2B | TRIDC1BG014960 |
| *T. aestivum* | A | *Bx6* | TraesCS2A01G051700 | chr2A | TraesCS1A01G085800 |
| *T. aestivum* | B | *Bx6* | TraesCS2B01G066000 | chr2B | TraesCS1B01G103700 |
| *T. aestivum* | D | *Bx6* | | | TraesCS1D01G087300 |
| *Z. mays* | | *Bx6* | Zm00008a014884 | chr04 | Zm00008a033413 |
| *D. oligosanthes* | | *Bx6* | | | OEL32741.1 |

| | | | | | |
|---|---|---|---|---|---|
| *E. haploclada* | | *Bx6* | eh_chr9.1288 | eh_chr9 | eh_chr6.2252 |
| *E. oryzicola* | AT | *Bx6* | AT09.1105 | AT09 | AT06.700 |
| *E. oryzicola* | BT | *Bx6* | BT09.1964 | BT09 | BT06.802 |
| | | | | | BT06.803 |
| *E. crus-galli* | AH | *Bx6* | AH09.1073 | AH09 | AH06.747 |
| *E. crus-galli* | BH | *Bx6* | | | BH06.788 |
| | | | | | BH06.790 |
| *E. crus-galli* | CH | *Bx6* | CH09.1289 | CH09 | CH06.821 |
| | | | | | CH06.823 |
| *E. colona* | DL1 | *Bx6* | | | DL06.2287 |
| | | | | | DL06.2288 |
| *E. colona* | DL2 | *Bx6* | DL09.1179 | DL09 | DL06.2287 |
| | | | | | DL06.2288 |
| *E. colona* | EL | *Bx6* | EL09.1058 | EL09 | EL06.1910 |
| | | | EL09.1059 | | EL06.1911 |
| | | | | | EL06.1913 |
| *E. colona* | FL | *Bx6* | FL09.1015 | FL09 | FL06.479 |
| | | | | | FL06.480 |

#### Table S3. Topology tests of two hypothesis on transfer of Bx genes in Triticeae from Panicoideae

| Datasets | Alignment with MAFFT | | | Alignment with MAFFT and Gblocks trimming | | |
|---|---|---|---|---|---|---|
| Topologies tested | PO vs Non-PO | | | PO vs Non-PO | | |
| | AU | RELL | SH | AU | RELL | SH |
| *Bx1* | 1.0000/4.98e-34 | 1.0000/0.0000 | 1.0000/0.0000 | 0.9999/5.41e-5 | 1.0000/0.0000 | 1.0000/0.0000 |
| *Bx2* | 1.0000/9.72e-9 | 1.0000/0.0000 | 1.0000/0.0000 | 0.9996/0.0004 | 1.0000/0.0000 | 1.0000/0.0000 |
| *Bx3/4/5* | 1.0000/1.43e-5 | 1.0000/0.0000 | 1.0000/0.0000 | 0.9981/0.0019 | 1.0000/0.0000 | 1.0000/0.0000 |
| *Bx8* | 1.0000/2.58e-44 | 1.0000/0.0000 | 1.0000/0.0000 | 1.0000/5.75e-5 | 1.0000/0.0000 | 1.0000/0.0000 |
| *Bx6* | 1.0000/1.18e-6 | 1.0000/0.0000 | 1.0000/0.0000 | 1.0000/4.4e-61 | 1.0000/0.0000 | 1.0000/0.0000 |

PO, Panicoideae origin; non-PO, non-Panicoideae origin; AU, approximately unbiased test; RELL, resampling estimated log-likelihood method; SH, Shimodaira-Hasegawa test.

```
                         *        20         *        40         *        60         *
N1 : MAFA-LKASPYLSSSSAAASS------SSSPALLPLPGQHAS----------ARVSFRPQAALTAPLAMGQA :  55
N2 : MAFTTMKASP----MSASSSS------------APVLRRCVAQPAR---VAAARRLAAAAASVALEASPVPA :  53
N3 : MAYA-VKASP-----SSSAPS------------LPFPRRRAAGAVV---TAGRRVKVRALAAAAALDPAAPA :  51
N4 : MAFA-LKASPSSTSSSSSAPSNHPRLRSAAAMAMPTPGRPASRAVA---AASAAASLEPAVVTPSPDSYWRA :  68
N5 : MAF-ALKAS--PSTSSSSLAVQ-----------SQLPRRAAAVATM-----PARRRAAAVRVMAIAAAPPAA :  53
N6 : MAF-AIKAASTSYS-TFSPADQ------------PSLSRLAATAAVKMPAGRGKAAAAVVIRAVAAAAAPLS :  58
N7 : MALFAVQAASTSQSSSSSPAVLQ---------QSPLPSRRAAAAAVKKMPQR-KKAAAVVRAVAAPPPAPPV :  62
N8 : MAS-AIKAASTSSQWSSSPAAVH---------SSPLSKRLPAAVA---MPGR-RRSVATVRAVAAVAPGAPA :  58
B1 : ---------------------------------------------------------------------- :   -
B2 : MAF-ALNAS--CYPSSFS---------------SLLPWRMAAAVMI-----PRRRNVLPV--IKAVAVAPPA :  47
B3 : MAF-ALKTS--FSASSFQAGPSS----------SSLLPRR---------------RNGVSV--IRAVATVSTS :  43
B4 : MAF-SLNAS--SSPSTFQAAQS-----------SLLPRRLSAVVTK-----PRPRNVFAV--IRAVATVS-- :  49
     ma      a                           r

             80         *       100         *       120         *       140
N1 : ---AAAAAAEKCN-VSQTFSRLRKQGKTAFIPYITAGDPDLATTAAALKLLDYCGADVIELGLPYSDPIADG : 123
N2 : A---AAAAVERRMSVSQTMSKLKEKGKTAFIPYITAGDPDMGTTAEALRLLDACGADVIELGVPFSDPYADG : 122
N3 : P---AAAGKRGRLSVSQTLSSLREKGKTAFIPYITAGDPDLETTAEALRLLDACGADVIELGMPFSDPYADG : 120
N4 : PLPLTPAASERIPTVAQAMSRARAHGKTAFIPYITAGDPDLGTTAEALRLLDALGADVIELGMPFSDPSADG : 140
N5 : PAPARPAGGRCL-PVSQTFAKLKAKGKTAFIPYITAGDPDLATTAEALRVLDACGADVIELGVPFSDPYADG : 124
N6 : PAPARSAGKRC-LPVSQTMARLKAQGKTAFIPYITAGDPDLATTAEALRLLDACGADVIELGVPFSDPYADG : 129
N7 : PGPPKPAGERCRLPVSQTMSRLKAQGKTAFIPYITAGDPDLATTAEALRLLDACGADVIELGVPFSDPYADG : 134
N8 : KLTAG-AGGRC-LPVSQTMSRLRAQGKTAFIPYITAGDPDLATTAEALRLLDACGADVIELGVPFSDPYADG : 128
B1 : ------------------MAALMAKGKTAFIPYITAGDPDLATTAEALRLLDGCGADVIELGVPCSDPYIDG :  54
B2 : PAPAKPAAVRSR-PVSVTMAKLMAKGKTALIPYITAGDPDLATTAEALRLLDACGADVIELGVPCSDPYVDG : 118
B3 : PAPAKPAAVTTTLPVSETLAKLMAKGKTAFIPYLTAGDPNLATTAEALRLLDACGADVIELGVPYSDPYLDG : 115
B4 : -APAKPAVRCR--PVSDTLAKLMAKGKVVYCTLYVA---------------HVYGS-TLELCLT-------- :  94
           a       vs t    l   GKta ipy tAgdp    ttaeal  ld cGadv6ELg6p sdp  dg

              *       160         *       180         *       200         *
N1 : PVIQASASRALAKG--ATLDTTMAMLKEVVPHLSCPVVLFTYYNPIMRRGVANFTAAAKEAGAHGLVVPDLP : 193
N2 : PVIQASASRALAAG--ATPEAVLSMLKEVTPELSCPVVLLSYLGPILRRGAANFTAAAKEAGVQGLIVPDLP : 192
N3 : PVIQASAARALAGG--ATADGVMSMLKEVTPELSCPVVLFSYFGPIVRRGVASVTAAVKEAGVQGLIVPDLP : 190
N4 : AVIQASAKRALAAG--ATTDAVMAMLREVTPELSCPVVIFSYFNPIVRRGTRSFAAAAREAGVKGLIIPDLP : 210
N5 : PVIQASSARALASG--TTPDAVLEMLREVTPELSCPVLLFSYFKPILCRGISDFTAAAKEAGIHGLLVPDLP : 194
N6 : PVIQASTARALASG--TTPDGVLAMLKEVTPELSCPVVLFSYFNPIVHRGLADFAAAAKEAGAHGLIVPDLP : 199
N7 : PVIQASMARARTTGGGATPDGVLAMLREVTPELSCPVVLFSYFNPIVRWGLPGFAAAVKDAGAHGLIVPDLP : 206
N8 : PVIQASAARALASG--TTPDGVLAMLKEVTPELSCPVVLFSYFNPIVRWGLADFAAAVKEAGVHGLIVPDLP : 198
B1 : PIIQASVARALASG--TTMDAVLEMLREVTPELSCPVVLLSYYKPIMSRSL----AEMKEAGVHGLIVPDLP : 120
B2 : PIIQASSARALAGG--ATMDGVLAMLKEVTPELSCPVVLFSYYRPILCRGL----AEIKEAGVHGLIVPDLP : 184
B3 : PIIQASTARALAGG--TTMDGVLEMLKEVTPELSCPVVLFSYYKPIFCQGL----AEIKEAGVHGLLVPDLP : 181
B4 : PSLQASSARALAGG--TTMDGVLAMLKEVTPELSCPVVLFSYYKPILFKGL----AEIKEAGVQGLLVPDLP : 160
     p 6QAS   RAla G    T d v6 ML4EVtPeLSCPV66f3Y  PI    g     A  4eAG  GL66PDLP

      220         *       240         *       260         *       280
N1 : LENAGIL--RNESIKNKLELVQLTTPVTPTGRMVQITQAAQGFVYLVSVVGVTGARSNVNPRVQHLLQEIKQ : 263
N2 : YVDTCTF--RSEAIKSNLELVLLTTPATPGERMKIITEASGGFVYLVSVNGVTGPRPKVNTRVEHLLQDIKL : 262
N3 : YVETCEF--RNESIKNNLELVLLTTPATPADRMKEITKASEGFVYLVSVNGVTGPRANVNPRVKDLLQEIRQ : 260
N4 : YDEIRAF--RKEAIQNSLELVLLTTPATPAERMKEIAKASEGFVYLVSVVGVTGARANVNPRVKDLLKEIRQ : 280
N5 : CVAECTL--KSEAMKNNLELVLLTTPATPEERMKEITKASEGFIYLVSVNGVTGPRANLNSRVQSLIQEVKQ : 264
N6 : YGATCAL--RSEATKNELELVLLTTPSTPEDRMKEIKEASEGFVYLVSISGVTGPRANVNTRVESLIKEVKQ : 269
N7 : YADTCAL--RSEAIKNDLELVLLTTPATPEERMKEITEASEGFVYLVSVNGVTGSRADVNTRVESLIQEVKQ : 276
N8 : YGNSCALTLRTEAIKNSLELVLLTTPSTPADRMGEITRASRGFVYLATVNGVTGPRANVNTRVQSLIQEVKQ : 270
B1 : YVAAHSL--WSEAKNNNLELVLLTTPAIPEDRMKEITKASEGFVYLVSVNGVTGPRANVNPRVESLIQEVKK : 190
B2 : YVAAHAL--WSEAKKNNLELVLLTTPAIPEERMKEITKASEGFIYLVTVNGVTGPRENVNLRVESLIQEIKK : 254
B3 : YVAAHAM--WSEAKKNNLELVLLTTPTTPEERMREITKASEGFVYLVSVDGVTGPRENVNSRVQSLIQEVKK : 251
B4 : YVAAHAL--WSDAKNNNLELVLLTTPTTPEERMKEITKASEGFIYLVSVNGVTGPRENVNSHVQSLIQEIKK : 230
     y           ea   n LELVlLTTP tP   RM   It As GF6YLv36 GVTG R   6N rV   L6qe64

      *       300         *       320         *       340         *       360
N1 : VTDKAVAVGFGISTPEHVKQVAQWGADGVIIGSAMVRQLGEASSPQE-----------------GLRR---- : 314
N2 : VTDKAVCVGFGISTPDHVRQIAGWGADGVIIGSAMVRQLGEAASPKQ-----------------GLKR---- : 313
N3 : VTDKAVAVGFGISTPDHVRQIAEWGADGVIIGSAMVRQLGEAASPKE-----------------GLRR---- : 311
N4 : VTDKEIAVGFGISTPDHVSQISEWGADGVIIGSAMVRQLGEADSPRE-----------------GLKR---- : 331
N5 : VTDKPVAVGFGISKPEHVKQIAEWGADGVIIGSAMVKQLERLALSRHRGPTSRHARTRMLYSKFGVRRNLFI : 336
N6 : VTDKPVAVGFGISKPEHVKQIAEWGADGVIIGSAMVRQLGEAGSPKE-----------------GLKR---- : 320
N7 : VTDKPVAVGFGISKPEHVKQIAEWGADGVIIGSAMVKQLGEATSPKE-----------------GLKR---- : 327
N8 : VTDIPVAVGFGISKPEHVKQIAEWGADGVIIGSAMVRQLGEAASPKE-----------------GLKR---- : 321
B1 : VTNKPVAVGFGISKPEHVK------------------QLGEAASPKQ-----------------GLRR---- : 223
B2 : VTDKPVAVGFGISKPEHVKQIARWGADGVIIGSAMVRQLGEAASPRE-----------------GLKR---- : 305
B3 : VTDKPVAVGFGISKPEHVKQIAQWGAHGVIIGSAMVRQLGEAASPKE-----------------GLKR---- : 302
B4 : VTTKPVAVGFGISKPEHVKQIAQWGADGVIIGSAMVRQLGEAPSPKE-----------------GLKR---- : 281
     VT  k 6aVGFGIS P HV q a wgadgviigsamv QLgea sp                 G64R
```

**Figure S1** Alignment of amino acid sequences of *Bx1* gene, related to Fig. 2c. See detailed sequences ID in Fig. 2a.

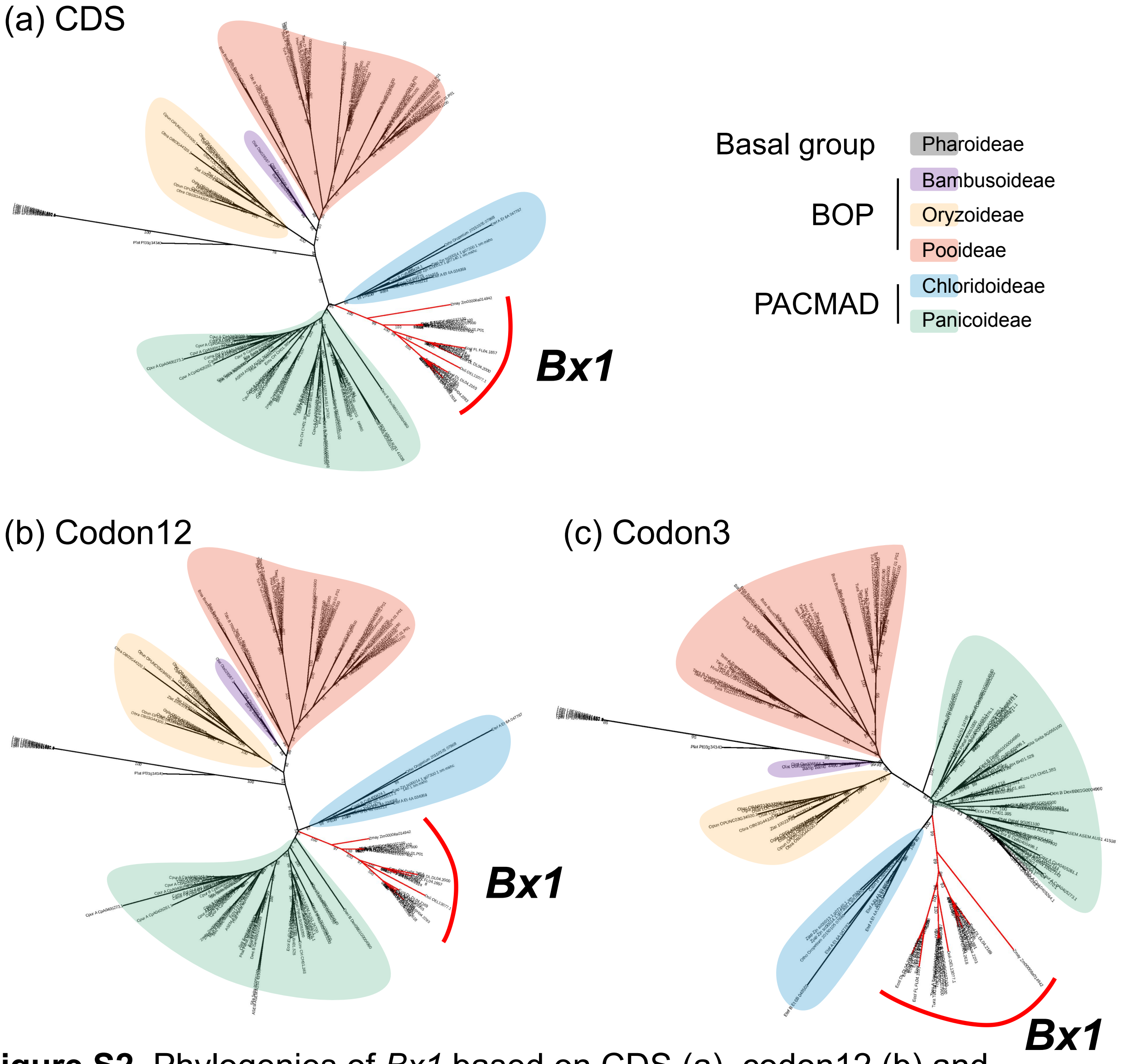


**Figure S2** Phylogenies of *Bx1* based on CDS (a), codon12 (b) and codon3 (c) datasets, related to Fig. 2a.

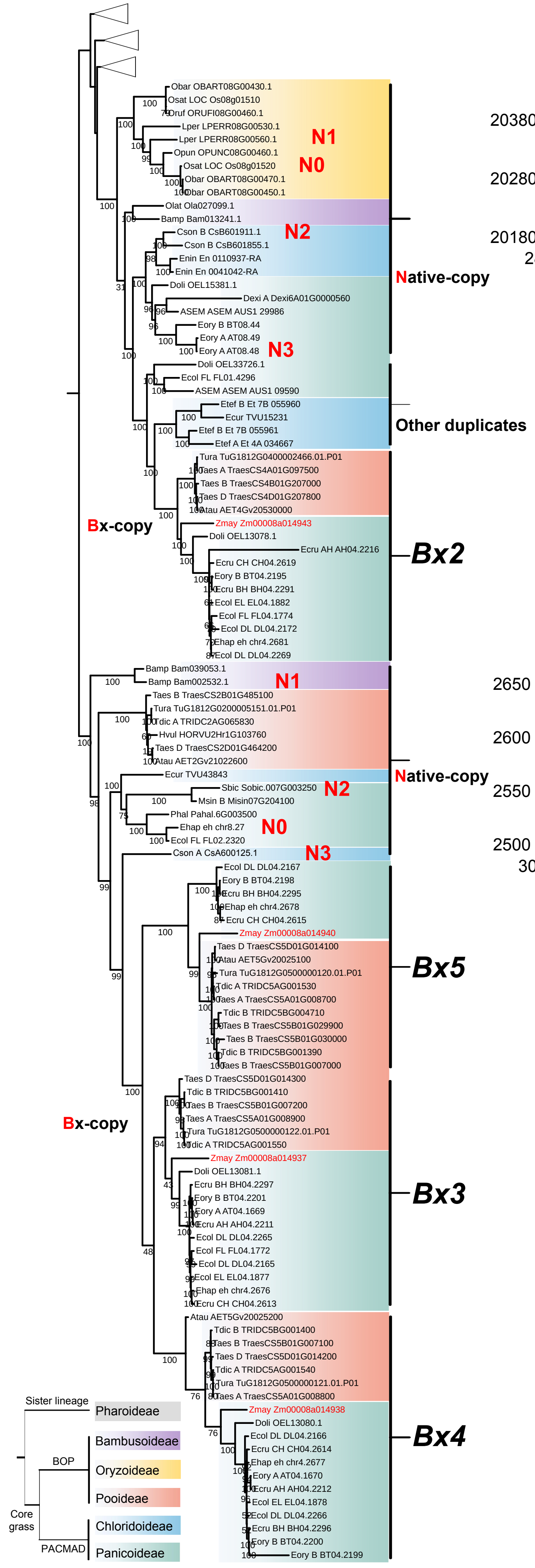


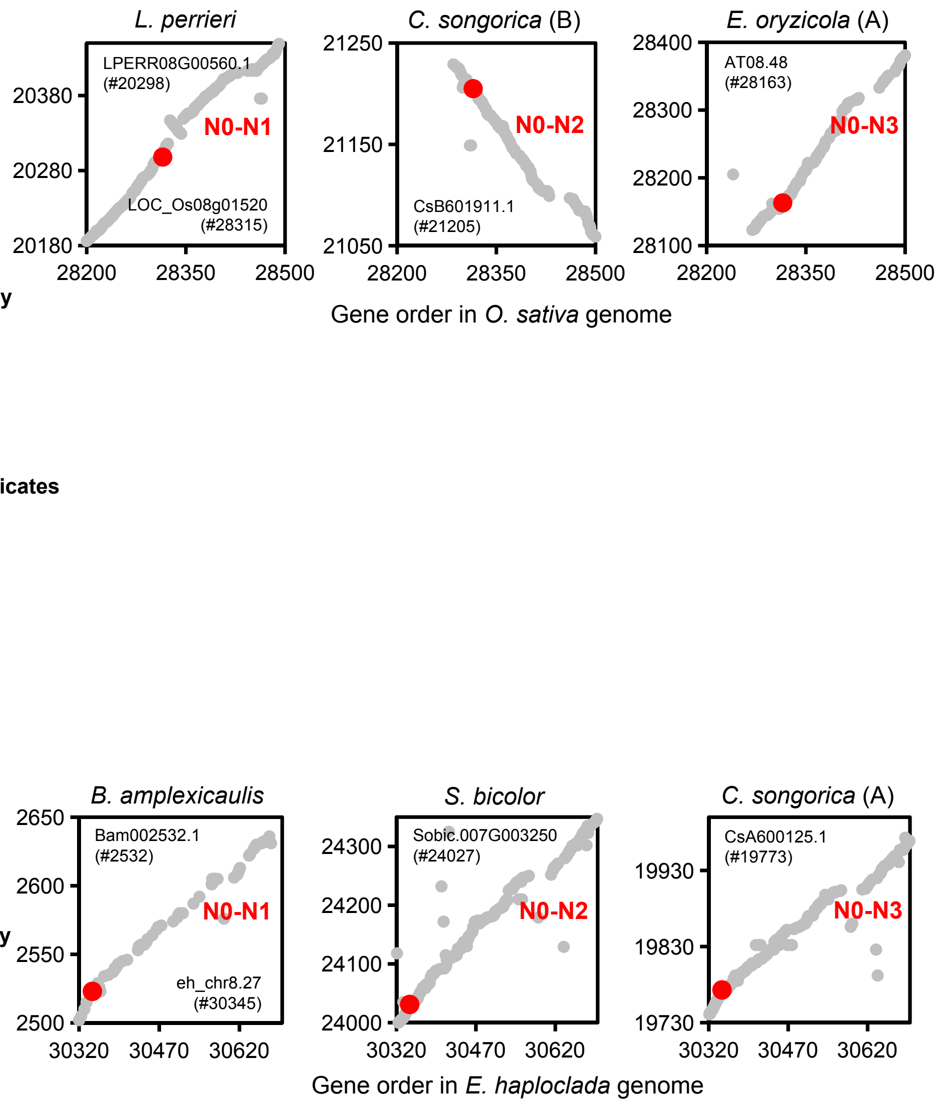


**Figure S3** Phylogeny of *Bx2* to *Bx5* and genomic synteny of *Bx2* and *Bx5* regions across the grass family

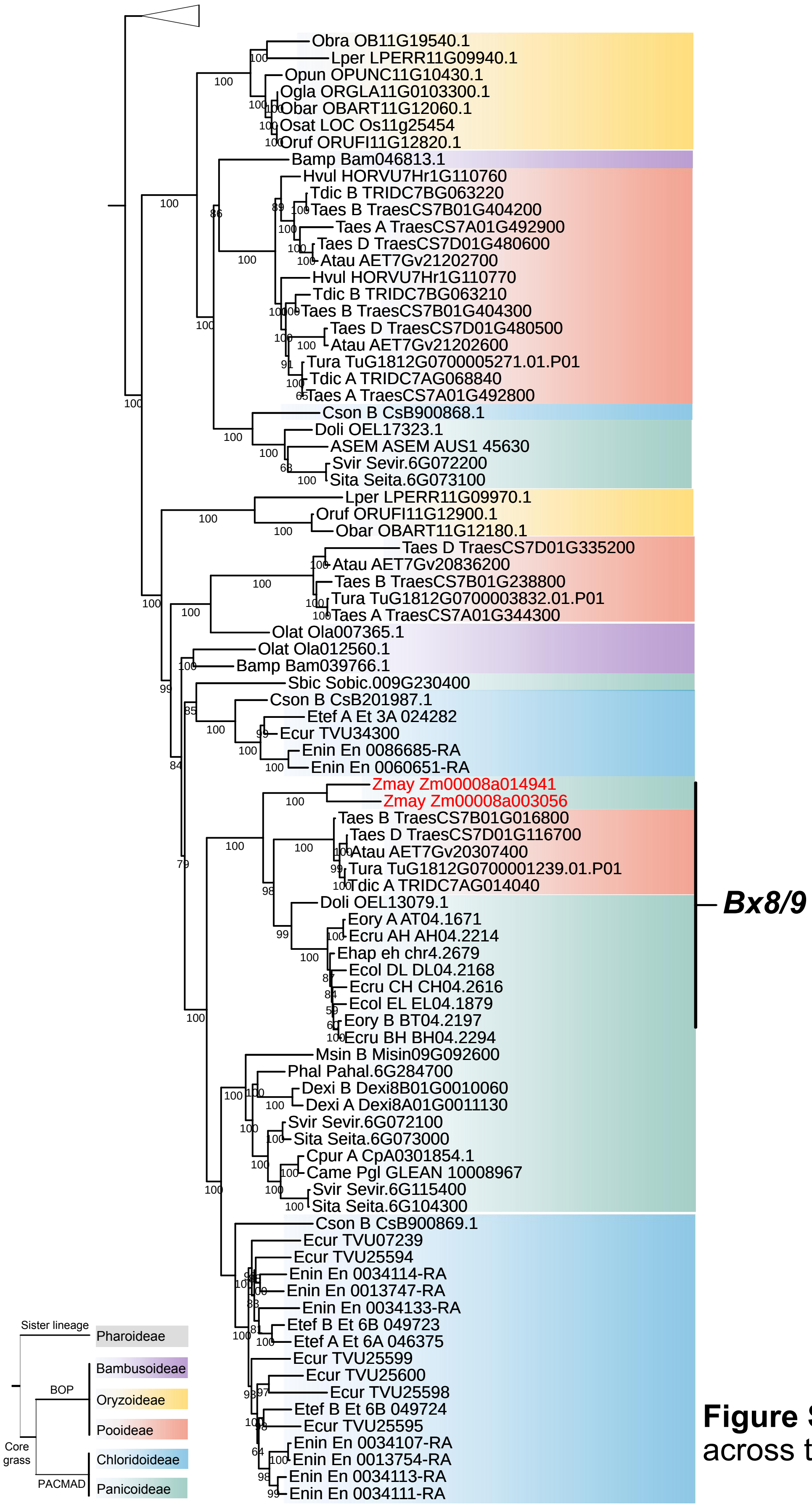


**Figure S4** Phylogeny of *Bx8* and *Bx9* across the grass family

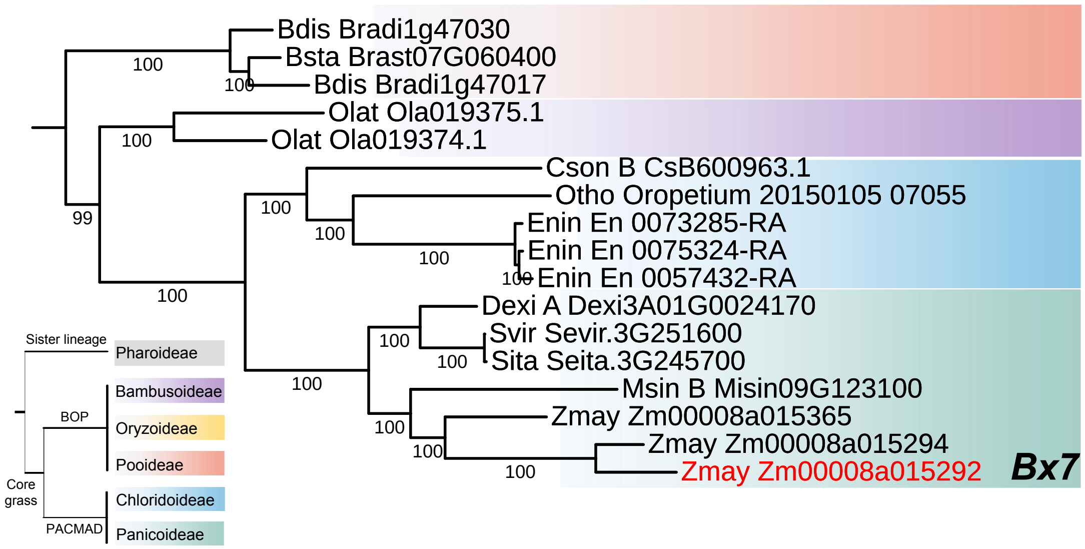


**Figure S5** Phylogeny of *Bx7* across the grass family

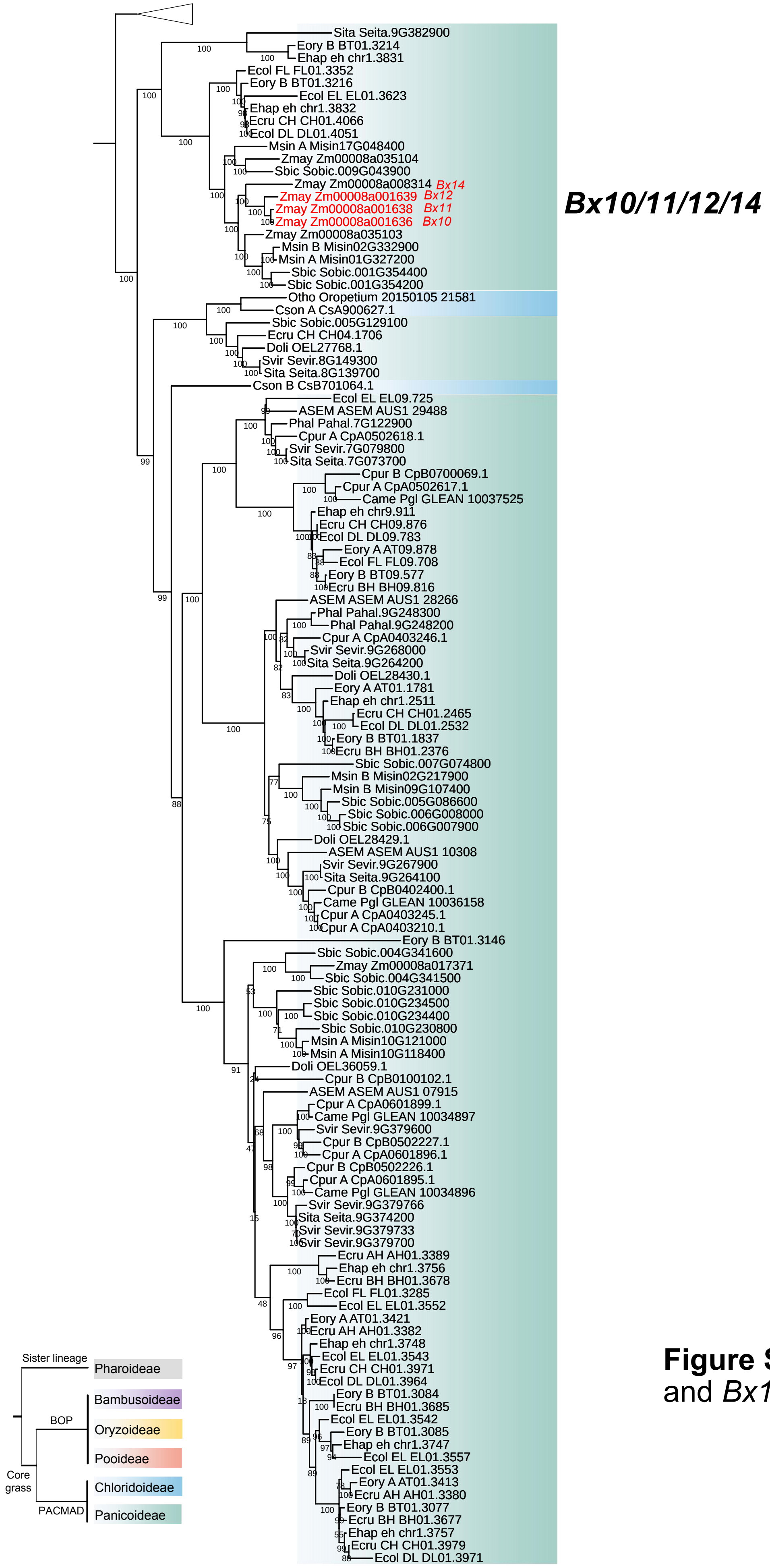


**Figure S6** Phylogeny of *Bx10* to *Bx12* and *Bx14* across the grass family

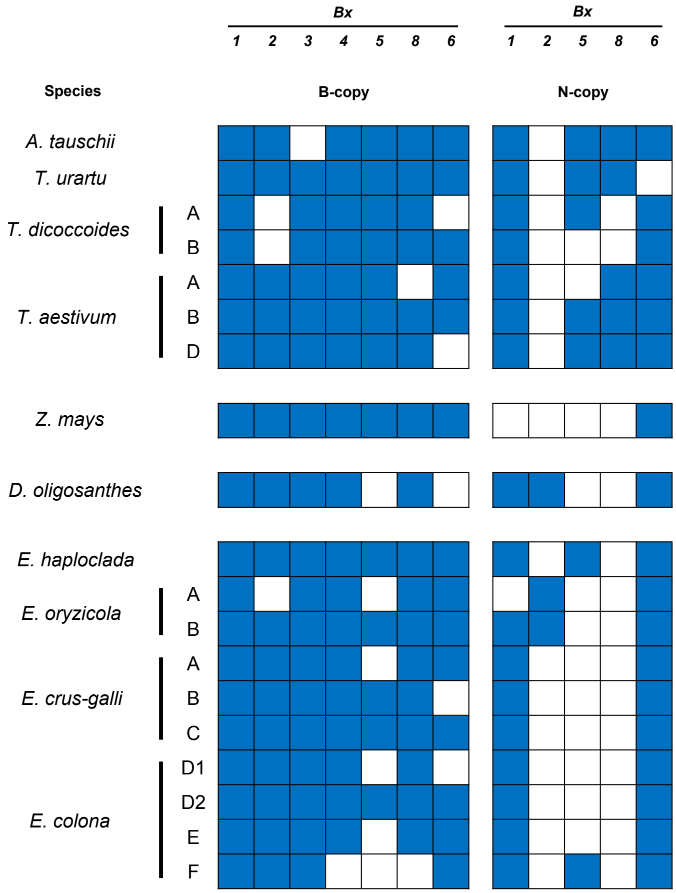


**Figure S7** Presence and absence of Bx genes (B-copy) and native analogues (N-copy). Blue grids represent presence and white represent absence.

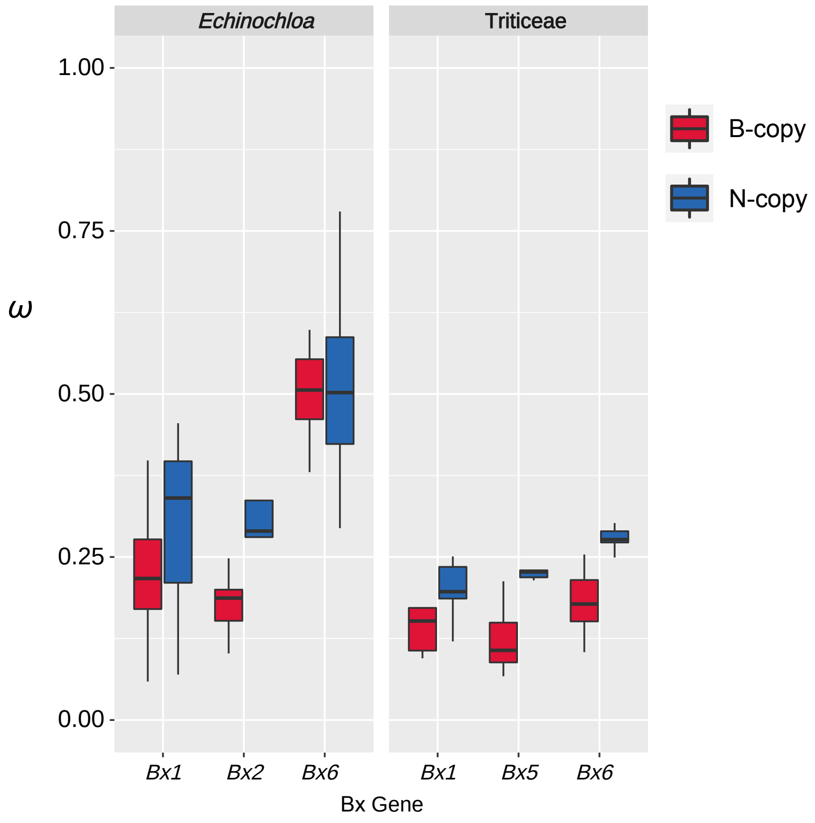


**Figure S8** Selection pressure of Bx genes in *Echinochloa* and Triticeae. In the box plots the horizontal line shows the median value, and the whiskers show the 25% and 75% quartile values of *ω*. B-copy, Bx genes; N-copy, native analogues of Bx genes.

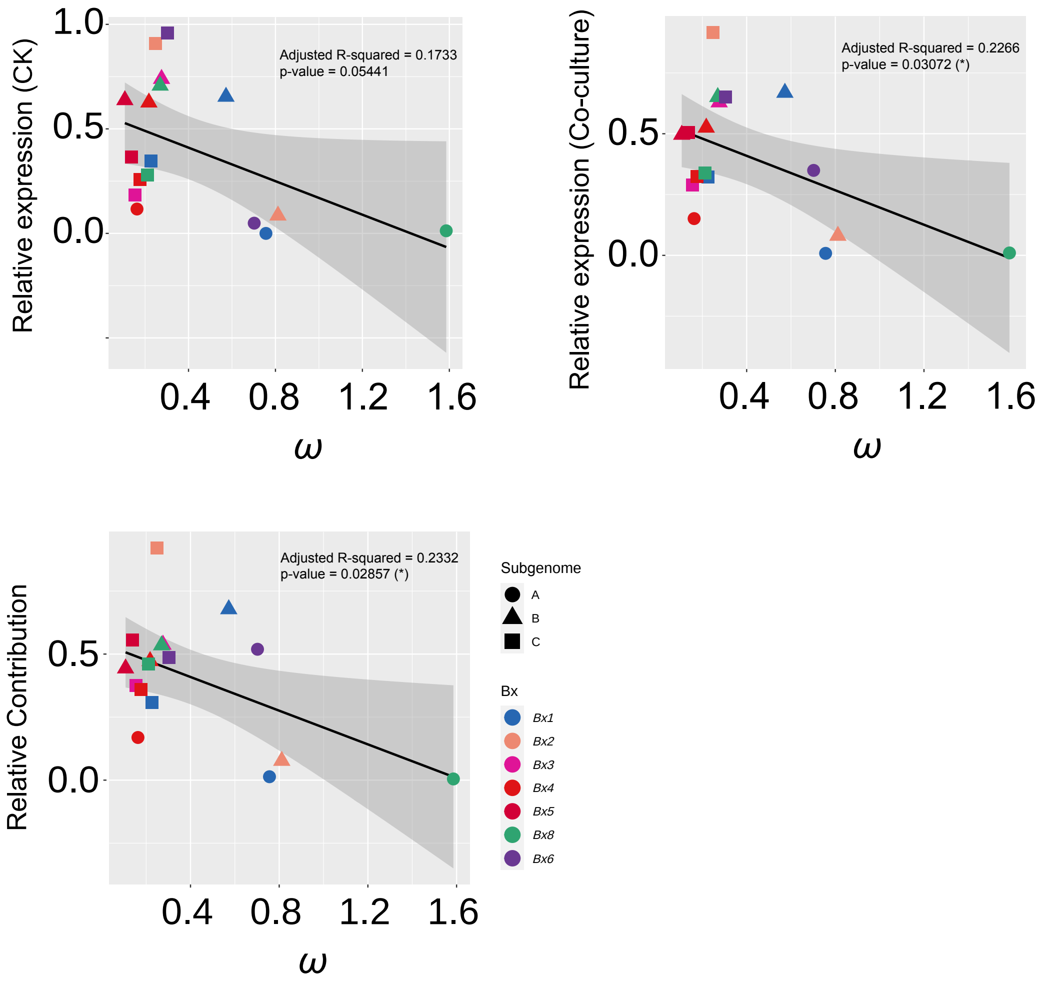


**Figure S9** Negative relationship between selection indicator *ω* values and expression or response dominance
